# The mitochondrial mRNA stabilizing protein, SLIRP, regulates skeletal muscle mitochondrial structure and respiration by exercise-recoverable mechanisms

**DOI:** 10.1101/2023.10.30.564600

**Authors:** Tang Cam Phung Pham, Steffen Henning Raun, Essi Havula, Carlos Henriquez-Olguín, Diana Rubalcava-Gracia, Emma Frank, Andreas Mæchel Fritzen, Paulo R. Jannig, Nicoline Resen Andersen, Rikke Kruse, Mona Sadek Ali, Jens Frey Halling, Stine Ringholm, Elise J. Needham, Solvejg Hansen, Anders Krogh Lemminger, Peter Schjerling, Maria Houborg Petersen, Martin Eisemann de Almeida, Thomas Elbenhardt Jensen, Bente Kiens, Morten Hostrup, Steen Larsen, Niels Ørtenblad, Kurt Højlund, Michael Kjær, Jorge L. Ruas, Aleksandra Trifunovic, Jørgen Frank Pind Wojtaszewski, Joachim Nielsen, Klaus Qvortrup, Henriette Pilegaard, Erik Arne Richter, Lykke Sylow

**Affiliations:** Department of Nutrition, Exercise and Sports, Faculty of Science, University of Copenhagen, Copenhagen, Denmark; Department of Biomedical Sciences, Faculty of Health and Medical Sciences, University of Copenhagen, Copenhagen, Denmark; Stem Cells and Metabolism Research Program, Faculty of Medicine, University of Helsinki, Finland; Division of Molecular Metabolism, Department of Medical Biochemistry and Biophysics, Karolinska Institutet, Stockholm, Sweden; Molecular and Cellular Exercise Physiology, Department of Physiology and Pharmacology, Karolinska Institutet, SE-17177 Stockholm, Sweden; Steno Diabetes Center Odense, Odense University Hospital, Odense, Denmark; Department of Biology, University of Copenhagen, Copenhagen, Denmark; British Heart Foundation Cardiovascular Epidemiology Unit, Department of Public Health and Primary Care, University of Cambridge, Cambridge, UK; Victor Phillip Dahdaleh Heart and Lung Research Institute, University of Cambridge, Cambridge, UK; Institute of Sports Medicine Copenhagen, Department of Orthopaedic Surgery M, Bispebjerg Hospital, Copenhagen, Denmark; Center for Healthy Aging, Faculty of Health and Medical Sciences, University of Copenhagen, Copenhagen, Denmark; Department of Sports Science and Clinical Biomechanics, University of Southern Denmark, Odense, Denmark; Clinical Research Centre, Medical University of Bialystok, Bialystok, Poland; Department of Clinical Research, University of Southern Denmark, Odense, Denmark; Institute for Mitochondrial Diseases and Aging, Cologne Excellence Cluster on Cellular Stress Responses in Aging-Associated Diseases (CECAD) and Center for Molecular Medicine (CMMC), Medical Faculty, University of Cologne, D-50931 Cologne, Germany

**Keywords:** SRA stem-loop interacting RNA-binding protein, mitochondrial dysfunction, skeletal muscle, exercise training, Drosophila

## Abstract

Decline in mitochondrial function associates with decreased muscle mass and strength in multiple conditions, including sarcopenia and type 2 diabetes. Optimal treatment could include improving mitochondrial function, however, there are limited and equivocal data regarding the molecular cues controlling muscle mitochondrial plasticity. Here we uncover the mitochondrial-mRNA-stabilizing protein SLIRP, in complex with LRPPRC, as a PGC-1α target that regulates mitochondrial structure, respiration, and mitochondrially-encoded-mRNA pools in skeletal muscle. Exercise training effectively counteracted mitochondrial defects induced by loss of LRPPRC/SLIRP, despite sustained low mitochondrially-encoded-mRNA pools, via increased mitoribosome translation capacity. In humans, exercise training robustly increased muscle SLIRP and LRPPRC protein content across exercise modalities and sexes, yet this increase was less prominent in subjects with type 2 diabetes. Our work identifies a mechanism of post-transcriptional mitochondrial regulation in skeletal muscle through mitochondrial mRNA stabilization. It emphasizes exercise as an effective approach to alleviate mitochondrial defects by possibly increasing mitoribosome capacity.

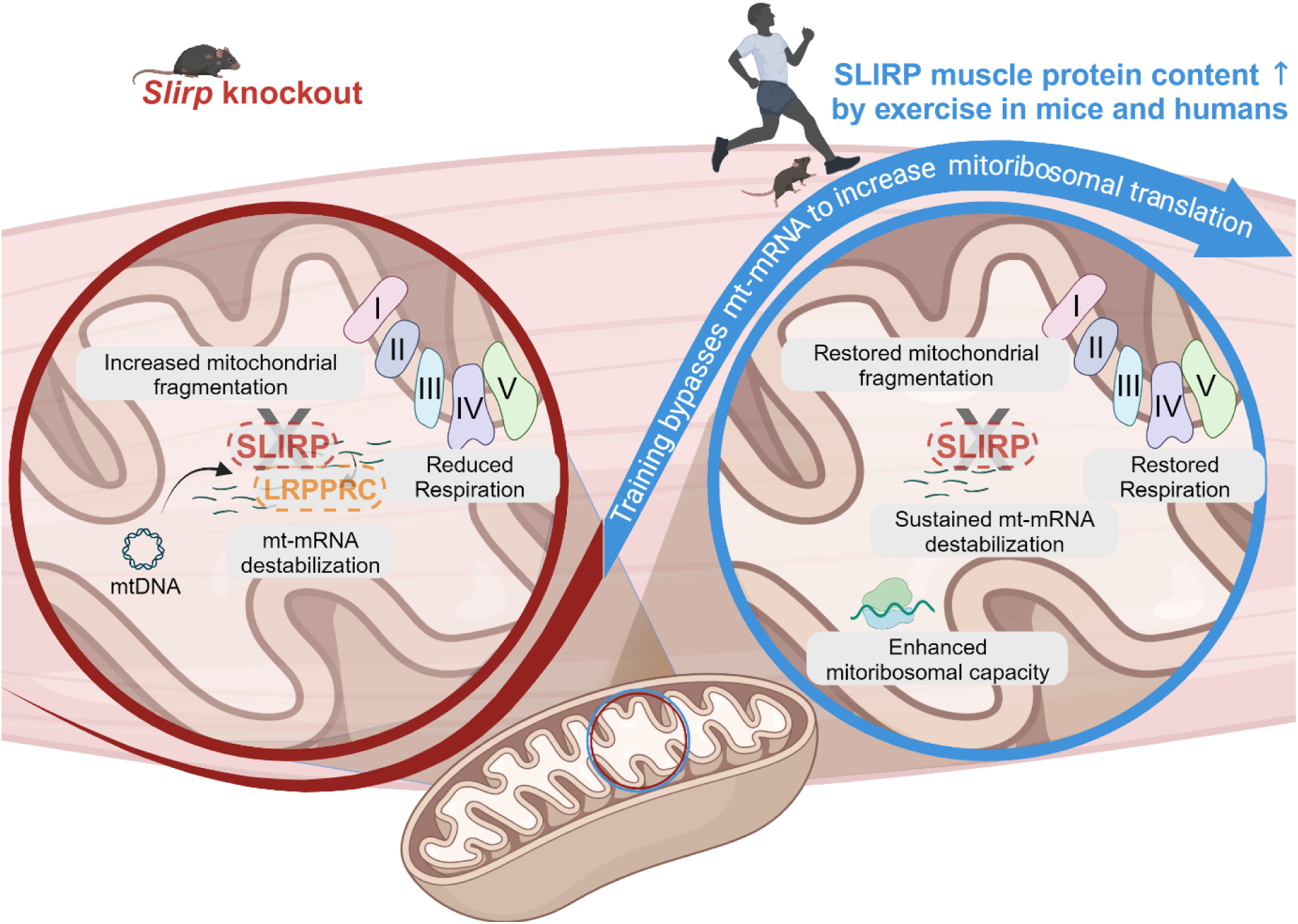

## Introduction

Mitochondrial homeostasis and function are vital for skeletal muscle physiology, primarily influenced by the ability to meet energy demands through oxidative phosphorylation (OXPHOS) in mitochondria^1^. The decline in mitochondrial function is associated with decreased muscle mass and strength in multiple conditions, including sarcopenia, type 2 diabetes (T2D), and cancer. Reduced muscle function adversely impacts health and quality of life^1–9^ and is associated with increased all-cause mortality. Mounting evidence suggests that improvements in mitochondrial metabolism contribute to the exercise-induced functional benefits, as mitochondrial preservation protects against the age-associated decline in skeletal muscle mass and performance^10,11^. The beneficial effects of exercise training are more potent than any drug in preserving muscle mass and function, positioning exercise training as a frontline strategy for the prevention and treatment of sarcopenia, T2D, and cancer^12–14^. Viewing mitochondrial control through the lens of exercise biology is a strategy to gain new mechanistic insights into mitochondrial regulation, which is currently poorly understood. With an aging population and no FDA/EMA-approved drugs to treat muscle functional decline, there is an urgent unmet need to identify potential treatment targets.

OXPHOS proteins are of dual origin and transcribed in spatially distinct cellular compartments. Nuclear (n)DNA encodes for the majority of OXPHOS proteins. Yet, 13 essential OXPHOS proteins are encoded by the small circular high-copy number mitochondrial DNA (mtDNA)^15,16^. In non-muscle systems, a protein complex comprised of steroid receptor RNA activator stem-loop interacting RNA-binding protein (SLIRP) and the leucine-rich pentatricopeptide repeat containing protein (LRPPRC) mediates mitochondrial-mRNA (mt-mRNA) stability and polyadenylation of most mtDNA-encoded OXPHOS transcripts^17–21^. Stabilization of mt-mRNA is critical for protein translation by the mitochondrial ribosome (mitoribosome). Yet, the mitoribosome comprises 82 mitoribosomal proteins encoded by nuclear genes, and 12S and 16S mt-rRNAs, highlighting the requirement for intricate coordination between the processes of cytosolic and mitochondrial protein synthesis^18,19^. Despite the high mitochondrial abundance in skeletal muscle, the complex interaction between LRPPRC/SLIRP-mediated posttranscriptional processes, mitoribosomal translation, and mitochondrial function has not been studied before in skeletal muscle. Elucidating the underlying mechanisms for stabilizing mt-mRNA could not only facilitate our understanding of the fundamental energy metabolism in muscle, but also aid intervention strategies for diseases associated with skeletal muscle mitochondrial defects.

Endurance exercise training potently increases mitochondrial mass in skeletal muscle, in part due to the increased content of nDNA- and mtDNA-encoded mitochondrial proteins^1,22^. Importantly, exercise training retains its ability to improve oxidative capacity not only in healthy individuals but also in patients with mtDNA mutations, with common PGC-1α-mediated mitochondrial adaptive responses shared between both groups^13^. These results suggest that exercise training induces adaptations to reinforce the mitochondria’s ability to efficiently respond to increased energy demands, even in the presence of mtDNA mutations, that may negatively affect mitochondrial transcription and translation. However, a notable knowledge gap persists in understanding the role of mitochondrial posttranscriptional processes, specifically in mt-mRNA stabilization and translation, in skeletal muscle biology and following exercise training. Yet, that knowledge is needed to gain a comprehensive understanding of the adaptive responses to exercise training.

Since SLIRP, a key player in mitochondrial posttranscriptional gene expression, is markedly upregulated in mouse skeletal muscle by exercise training^23^, we hypothesized that SLIRP would regulate mitochondrial function in skeletal muscle at rest and in response to exercise training. Our findings illuminate SLIRP’s role in regulating mt-mRNA transcript levels in skeletal muscle, downstream of PGC-1α1. Knockout (KO) of SLIRP led to damaged and fragmented mitochondria alongside lowered respiration. Intriguingly, exercise training could compensate for the absence of SLIRP, leading to improvements in mitochondrial integrity and respiratory capacity. Our findings imply the activation of complex exercise-induced molecular signaling in skeletal muscle which intriguingly bypasses mt-mRNA defects, possibly through enhanced mitoribosomal translation.

## Results

### *Slirp* knockout caused defects in mitochondrial structure and respiratory capacity in mouse muscle

Protein profiling of five different skeletal muscle tissues showed that SLIRP content was highest in the oxidative soleus compared to glycolytic extensor digitorum longus (EDL, Fig. 1A, Supplementary Fig. 1A), in accordance with the potential critical role for SLIRP in oxidation. Moreover, SLIRP was present across a diverse array of tissues in mice and was highly abundant in energy-demanding tissues such as liver, kidney, brown adipose tissue, heart, and skeletal muscle (Fig. 1A, Supplementary Fig. 1A), in line with previously reported *Slirp* mRNA levels^24^.

**Fig 1.**
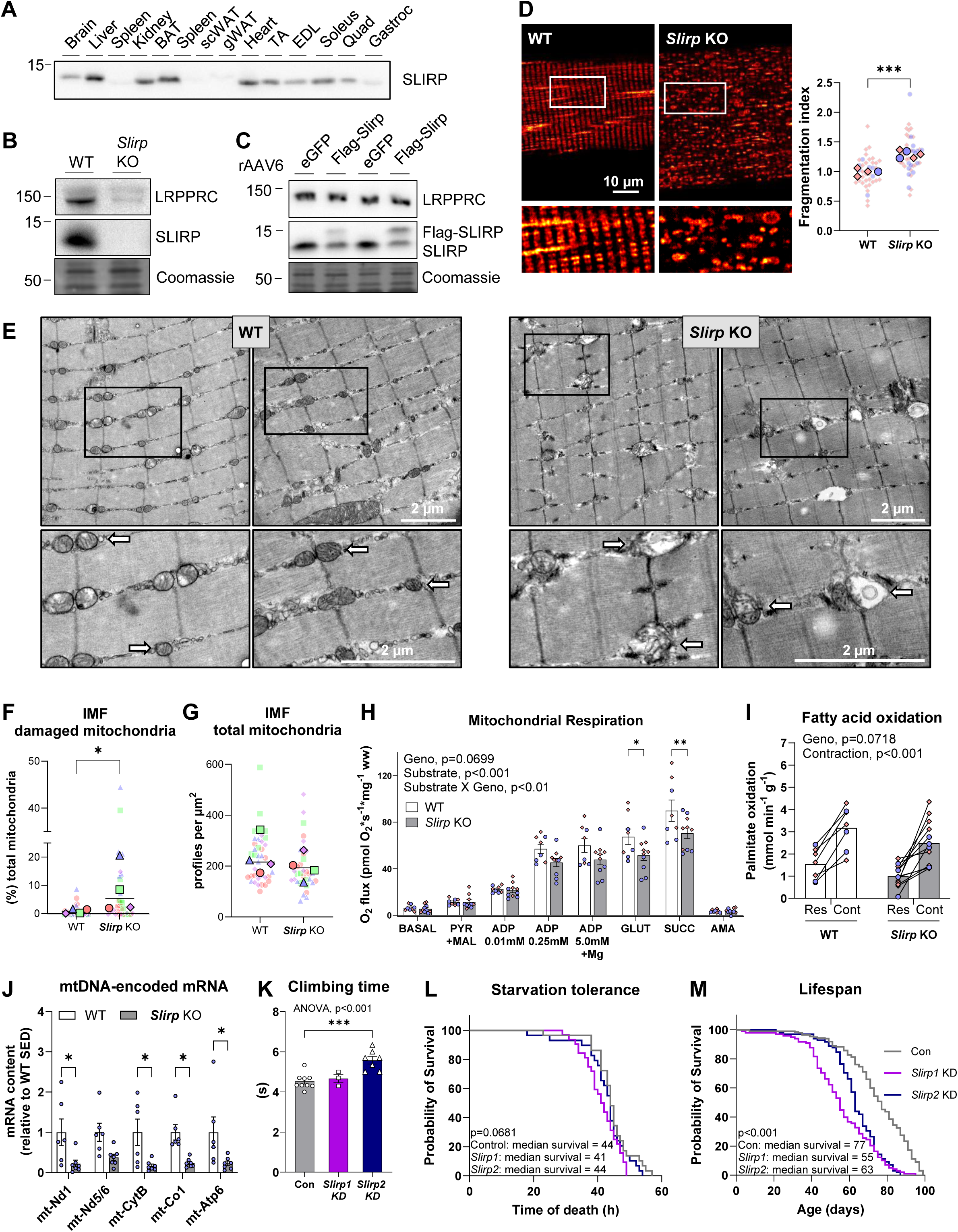
*Slirp* knockout led to defects in mitochondrial structure and function, and reduced lifespan. (A) SLIRP protein content across different wild-type tissues (n=4, female C57BL/6J, 12 weeks of age) assessed by immunoblotting. (B) SLIRP and LRPPRC protein content in tibialis anterior muscle of *Slirp* knockout (KO) and littermate wildtype (WT) mice, and recombinant adeno-associated virus serotype 6 encoding SLIRP (rAAV6:SLIRP) and rAAV6:eGFP as control mice. (C) SLIRP and LRPPRC protein content in tibialis anterior muscle of recombinant adeno-associated virus serotype 6 encoding SLIRP (rAAV6:SLIRP) and rAAV6:eGFP in the contralateral leg as control. (D) Confocal microscopy of mitochondrial network structure using a mitochondrial membrane potential probe (TMRE+) in flexor digitorum brevis muscle fibers of *Slirp* KO and WT mice (n=4-5, 7-10 fibers per mouse). Corresponding fragmentation index are presented as super plots^94^; small symbols represent each fiber, large symbols represent the mean of fibers from each mouse that are color- and symbol-coded for each sex (**○** (blue) male, ◇ (red) female). (E) Transmission electron microscopy images of *Slirp* KO and WT gastrocnemius muscle. (F and G) Quantification of percentage of damaged intramyofibrillar (IMF) mitochondria within total mitochondrial and relative volume of mitochondria. Small symbols refer to damaged or total mitochondria per fiber, large symbols of identical symbol refer to average of fibers per biological replicate (female *Slirp* KO and WT gastrocnemius muscle, n=4/group). (H) Mitochondrial respiration in of *Slirp* KO and WT gastrocnemius muscle (n=4-5/group, **○** (blue) male, ◇ (red) female). (I) Fatty acid oxidation in isolated *Slirp* KO and WT soleus muscle at rest and in response to contraction (n=3-6/group, **○** (blue) male, ◇ (red) female). (J) RT-qPCR analysis of mitochondrial transcript levels in gastrocnemius of male *Slirp* KO and WT mice (n=6-7/group). (K-M) Climbing time during climbing assay, starvation tolerance and life span of control (*Mef2-GAL4*>GD control 6000), *SLIRP1* (*Mef2-GAL4*>*SLIRP1* RNAi 51019 GD) and *SLIRP2* (*Mef2-GAL4*> *SLIRP2* RNAi 23675 GD) knockdown flies. Data are shown as mean ± SEM, including individual values, where applicable. Geno, main effect of genotype; substrate, main effect of substrate addition; Substrate X Geno, interaction between genotype and substrate; Contraction, main effect of contraction. *p<0.05, **p<0.01, ***p<0.001, as per Unpaired Student’s t test (D, J), Mann Whitney test (F, G), Two-way RM ANOVA with Šídák’s multiple comparisons test (H, I), ordinary one-way ANOVA with Dunnett’s multiple comparisons test (K), Log-rank (Mantel-Cox) test (L, M).

In skeletal muscle of global *Slirp* KO mice, SLIRP protein was undetectable. Moreover, SLIRP’s binding partner LRPPRC showed 90% reduction in protein content (Fig. 1B), indicative of their co-stabilization. To confirm co-stabilization, we administered intramuscular injections of recombinant adeno-associated viral serotype 6 (rAAV6):*Slirp*. As expected, the absence of concomitant upregulation of LRPPRC, failed to increase total SLIRP protein content due to downregulation of endogenous SLIRP protein (Fig. 1C). These findings show that SLIRP and LRPPRC stabilize each other in skeletal muscle, aligning with findings from non-muscle tissue^17,18,20^.

We next determined the role of SLIRP for mitochondrial structure in skeletal muscle, using the membrane potential probe, tetramethylrhodamine ethyl ester (TMRE+), in intact flexor digitorum brevis (FDB) muscle fibers of *Slirp* KO and littermate control wild-type (WT) mice. Interestingly, the finely interconnected mitochondrial network, obvious in WT fibers, was disrupted in *Slirp* KO muscle fibers, evident by a 30% increased mitochondrial fragmentation index (Fig. 1D). Transmission electron microscopy (TEM) analysis of gastrocnemius muscle provided further insights into the ultrastructural changes of mitochondrial morphology in *Slirp* KO (Fig. 1E). Half of the KO gastrocnemius muscles displayed reduced density of the matrix, disarray of the cristae, and substantial enlargement and vacuolation of intermyofibrillar (IMF) mitochondria (Fig. 1E). Quantitative analysis of the mitochondria showed that the percentage of damaged IMF mitochondria was increased (Fig. 1F).

However, there were no significant alterations in the total number of IMF mitochondria (Fig. 1G), number of damaged or the total number of subsarcolemmal mitochondria (Fig. 1G), nor any changes in number of damaged or total number of subsarcolemmal mitochondria (not shown). The percentage of damaged mitochondria in *Slirp* KO muscle varied between 2-20% of total mitochondria compared to <2% of damaged mitochondria in WT muscle (Fig. 1E, F). Other structural parameters of IMF mitochondria, such as mitochondrial area, aspect ratio, elongation, convexity, perimeter, sphericity, and diameter did not display any significant changes in *Slirp* KO muscles, likely due to heterogeneity in the extent of damage (Fig. 1F, Supplementary Fig. 1B-H).

In agreement with the observable abnormalities in mitochondrial structure, maximal respiratory capacity was reduced in permeabilized gastrocnemius muscle fibers of *Slirp* KO mice compared to WT (Fig. 1H). This was particularly evident when adding glutamate to assess maximal complex I linked respiratory activity (−24%) and succinate to assess complex I+II linked respiratory capacity (−21%). These results indicate an important role for SLIRP in skeletal muscle respiration, which contrast findings in isolated mitochondria from liver and heart tissues, where *Slirp* KO did not compromise mitochondrial respiration^18^. In agreement with lower respiration, fatty acid oxidation tended (p=0.0718) to be lower in in intact incubated soleus muscle of *Slirp* KO mice (Fig. 1I). However, electrically-induced muscle contraction increased fatty acid oxidation similarly in both genotypes, suggesting that fatty acid utilization for fuel during muscle contraction does not depend on SLIRP (Fig. 1I).

With the suggested role of SLIRP as an mt-mRNA stabilizing protein in non-muscle cells^18^, we determined mtDNA-encoded mRNA transcripts. Intriguingly, we observed a 60-80% reduction in *mt-Nd1*, *mt-Nd5/Nd6* (Complex I), *mt-CytB* (Complex III), *mt-Co1* (Complex IV), and *mt-Atp6* (ATP synthase) in gastrocnemius muscle (Fig. 1J), suggesting that SLIRP stabilizes mt-mRNA in skeletal muscle.

Together, these results suggest that mt-mRNA stabilization via SLIRP is required for proper mitochondrial network structure and morphology, and respiration in mouse muscle.

### SLIRP knockdown impaired locomotion and reduced lifespan in flies

To determine the long-term consequences of SLIRP deficiency on the whole organism, we utilized the UAS-GAL4 system^25^ for muscle-specific (*Mef2*-GAL4>) knock-down (KD) of *SLIRP1* and *SLIRP2* in *Drosophila (D.) melanogaster*. Only one *SLIRP* gene is present in mouse and human, while two *SLIRP* genes exist in *D. melanogaster* (Flybase annotation symbols (http://flybase.org): CG33714 and CG8021) likely to have originated from gene duplication events^26^. This consequently gives rise to two fly orthologue proteins of the human and mouse SLIRP, denoted *SLIRP1* (CG33714) and *SLIRP2* (CG8021)^26^. To test physical functionality, we subjected the flies to a negative geotaxis (climbing) assay^27^ and found that *SLIRP2* KD, but not *SLIRP1* KD, flies climbed 23% slower than control flies, indicative of impaired muscle function (Fig. 1K). The detrimental effects of *SLIRP* KD in the muscle became further apparent following starvation. Median fasting survival of *SLIRP1* KD flies was 41 h, compared with 44 h for control and *SLIRP2* KD flies (Fig. 1L, p=0.0681). Muscle-specific *SLIRP* deficiency, irrespective of orthologue, had a deleterious effect on lifespan. While the median survival of control flies was 77 days, median survival of *SLIRP1* and *SLIRP2* KD flies was only 55 and 63 days, respectively, and thus both *SLIRP1* and *SLIRP2* KD significantly reduced lifespan (Fig. 1M).

Taken together these findings in the flies show that *SLIRP* KD-induced mitochondrial defects can have detrimental long-term consequences such as impaired muscle function, starvation intolerance, and reduced life span.

### Muscle SLIRP protein content increases in response to exercise training, under the regulation of PGC-1α1

Having established critical functional roles of SLIRP in skeletal muscle mitochondrial morphology and respiratory capacity with detrimental effects on lifespan, we next investigated the upstream regulation of SLIRP protein content. Our recent work pinpointed SLIRP as a protein responsive to exercise training^23^; yet the mechanisms governing its induction and functions remained elusive. Knowing that exercise potently increases PGC-1α protein and mRNA levels in both mouse and human skeletal muscle^28–31^, and that PGC-1α is an important regulator of mitochondrial biogenesis, respiration, and quality control^32^, we explored the regulation of SLIRP by PGC-1α. The *Ppargc1a* gene encodes several PGC-1α isoforms, including isoform 1 (PGC-1α1) and isoform 4 (PGC-1α4), with distinct regulation and biological functions^33,34^. Overexpression (OE) of PGC-1α1 in mouse skeletal muscle^33,35^, associating with endurance exercise training-like adaptations^33,35,36^, resulted in approximately 8.3-fold higher muscle SLIRP protein content, without a concomitant change in mRNA levels (Fig. 2A). On the contrary, PGC-1α4 OE^37^, inducing resistance-type exercise training adaptations such as muscle hypertrophy and strength^33^, had no effect on SLIRP protein content in mouse gastrocnemius muscle (Fig. 2B). This suggests that SLIRP is an unrecognized player in PGC-1α1-regulated oxidative metabolism.

**Fig 2.**
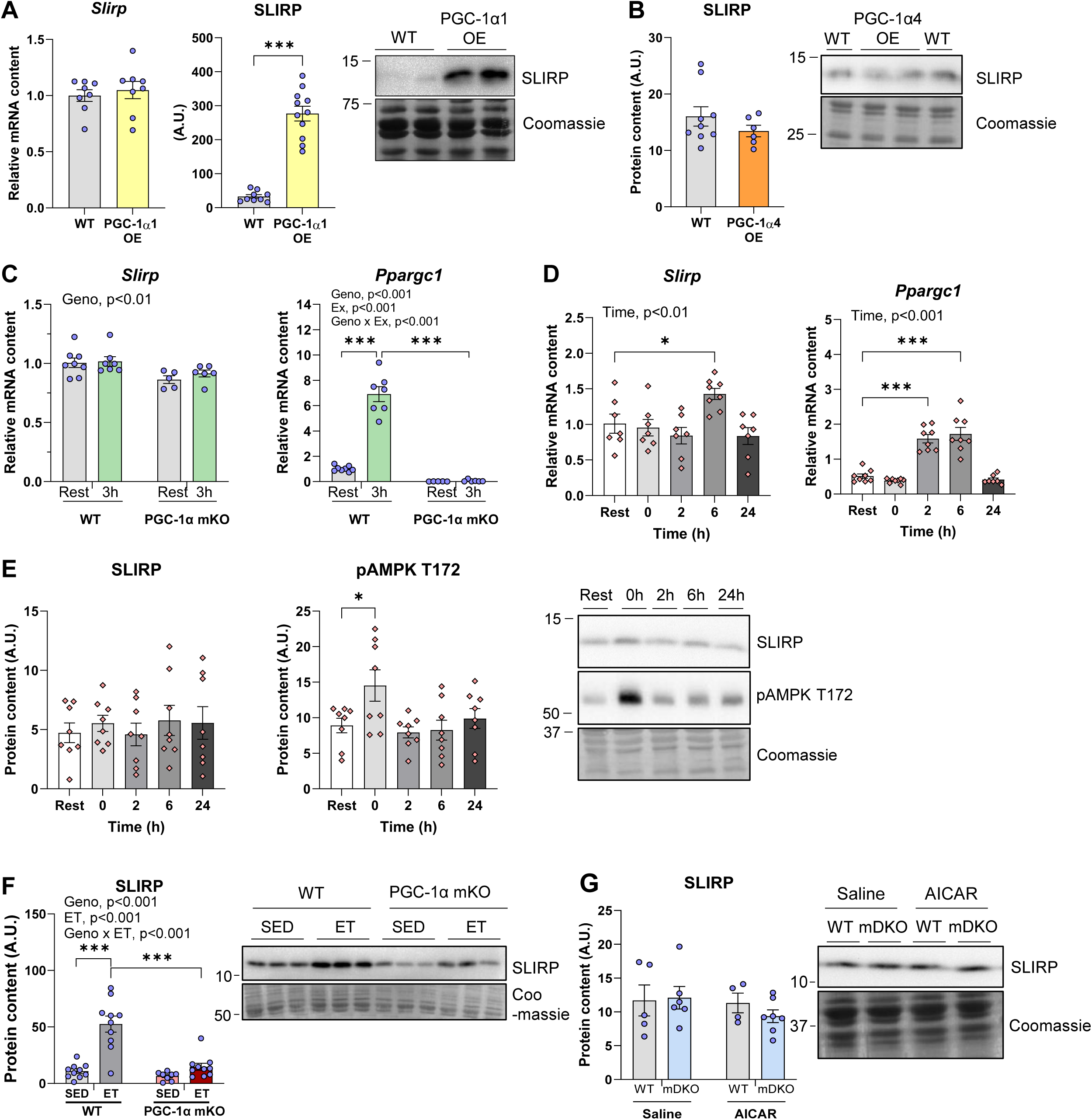
SLIRP is upregulated by ET and is a target of PGC-1α. (A) RT-qPCR analysis of *Slirp* transcript levels relative to *Actb* levels and Western blot analysis of SLIRP protein abundance in quadriceps with skeletal-muscle-specific transgenic expression of PGC-1α1 (α1 OE; n=8-11/group) (B) Western blot analysis of SLIRP protein abundance in gastrocnemius with skeletal-muscle-specific transgenic expression of PGC-1α4 (α4 OE; n=6-9/group). (C) RT-qPCR analysis of *Slirp* and *Ppargc1α* transcript levels relative to *Hprt* levels in gastrocnemius of muscle-specific PGC-1α knockout (PGC-1α mKO) and littermate control (WT) mice at rest and 3h after acute exercise bout (n=5-8/group). (D) RT-qPCR analysis of *Slirp* and *Ppargc1α* transcript levels relative to *Actb* levels in WT quadriceps at rest, immediately after, 2 h, 6 h or 24 h after acute exercise bout (n=8/group). (E) Western blot analysis of SLIRP and pAMPK T172 protein abundance in WT quadriceps at rest, immediately after, 2 h, 6 h or 24 h after acute exercise bout (n=8/group). (F) Western blot analysis of SLIRP protein abundance in quadriceps of sedentary (SED) and 12-week ET PGC-1α mKO and WT mice (n=9-10/group). (G) Western blot analysis of SLIRP protein abundance in quadriceps of muscle-specific AMPKα1 and -α2 double KO (mDKO) and WT mice treated with/without AICAR (n=4-7/group). Data are means ± SEM, including individual values. Geno, main effect of genotype; Ex, main effect of acute exercise; Geno x Ex, interaction between genotype and acute exercise. *p<0.05, ***p<0.001, as per Unpaired Student’s t test (A), Two-way ANOVA with Šídák’s multiple comparisons test (C, F, G), and ordinary one-way ANOVA with Dunnett’s multiple comparisons test (D, E).

To test whether PGC-1α regulates *Slirp* mRNA in skeletal muscle following exercise, we determined *Slirp* and *Ppargc1a* (Exon 3-5) mRNA levels in tibialis anterior (TA) muscle 3 h into recovery from one exercise bout in mice lacking PGC-1α in skeletal muscle (PGC-1α mKO) and WT control littermates. *Slirp* mRNA was not upregulated 3 h post-exercise (Fig. 2C), in contrast to the expected^31^ increase in *Ppargc1a* (+6.8-fold; Fig. 2C) in WT mice. Interestingly, *Slirp* mRNA levels were only modestly 14% reduced in PGC-1α KO skeletal muscle (Fig. 2C), in line with the dissociation between *Slirp* mRNA levels and SLIRP protein levels observed in PGC-1α1 OE skeletal muscle (Fig. 2A).

These findings align with reports describing only moderate correlations between mRNA and protein levels following exercise^38–40^. Yet, with the interpretative limitations of a single post-exercise time point, we aimed to acquire a more detailed record of the temporal changes in *Slirp* transcripts in recovery from exercise. We subjected WT mice to an acute exercise bout at 60% of their maximal running capacity and harvested quadriceps muscles at rest, immediately after, 2h, 6h, and 24h post-exercise. The phosphorylation of AMP-activated protein kinase (AMPK) at T172 served as a validation of elevated metabolic stress during the exercise bout (Fig. 2E). We found that *Slirp* mRNA levels were 1.4-fold elevated 6h into the recovery period in quadriceps muscle (Fig. 2D), a time-point not included in our prior cohort (Fig. 2C), and partially concurrent with *PGC-1α1* mRNA levels that were elevated 2h (+3.0-fold) and 6h post-exercise (+3.3-fold; Fig. 2D). Following a single bout of exercise, we observed no changes to SLIRP protein content (Fig. 2E). These findings indicate that SLIRP underlies PGC-1α1-regulated control following acute exercise.

Given that SLIRP might be regulated by PGC-1α, we next turned to investigate the long-term dependency of exercise training-induced regulation of SLIRP protein by PGC-1α. In contrast to acute exercise, exercise training increased SLIRP protein content 4.8-fold in WT mice (Fig. 2F), recapitulating our earlier observations in mice engaged in voluntary wheel running^23^. The effect of exercise training on SLIRP protein content was markedly blunted by 70% in mice lacking PGC-1α in muscle compared to trained WT mice (Fig. 2F). Expectedly, in WT mice, exercise training potently upregulated the steady-state levels of multiple OXPHOS proteins including SDHB, UQCRC2, MTCO1 in skeletal muscle (Supplementary Fig. 1I-K). In mice lacking PGC-1α in muscle, SDHB (Supplementary Fig. 1I) and MTCO1 protein (Supplementary Fig. 1K) was 50-60% decreased in SED mice. Moreover, the effect of exercise training on SDHB (Supplementary Fig. 1I) and UQCRC2 protein content (Supplementary Fig. 1J) was blunted by 30% and 65%, respectively, in PGC-1α mKO compared to WT mice. In contrast to PGC-1α, the exercise-sensitive metabolic sensor AMPK^41^ did not seem to regulate SLIRP expression, as SLIRP protein abundance was comparable to controls in muscle-specific double KO of AMPKα1 and AMPKα2 and WT mice^42,43^ treated with or without the AMPK-activator AICAR for 4 weeks (Fig. 2G).

PGC-1α has marked effects on muscle endurance, partially attributed to its pivotal role in mitochondrial biogenesis, and respiration^35,44^. Our results suggest SLIRP as an unrecognized downstream target of PGC-1a in such exercise adaptations and oxidative metabolism.

### SLIRP is dispensable for organismal adaptations to exercise training, yet, needed for improving blood glucose regulation after exercise training in male mice

Having identified SLIRP as an exercise training responsive protein downstream of PGC-1a, we next determined SLIRP’s mechanistic involvement in exercise training-mediated organismal adaptations. To this end, we conducted a 10-week voluntary wheel-running training study in *Slirp* KO and WT littermate mice of both sexes commencing at ∼18 weeks of age. The experimental design is shown in Fig. 3A.

**Fig. 3.**
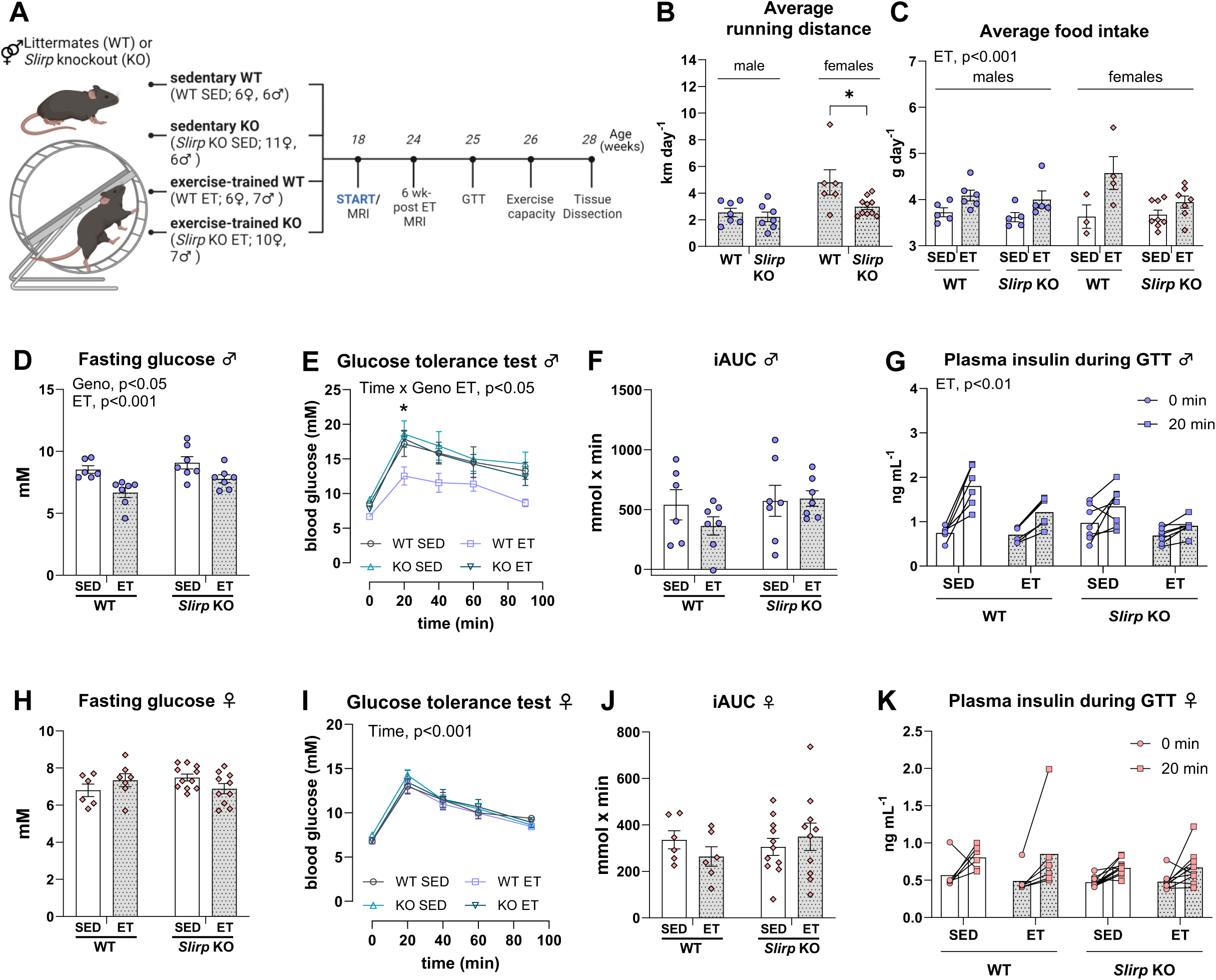
SLIRP is dispensable for organismal adaptations to ET, yet, needed for improving blood glucose regulation after ET in male mice. (A) Experimental design of 10-week exercise training (ET) intervention. (B) Average running distance of male and female ET *Slirp* KO or littermate control (WT) mice measured for 6 weeks (n=6-11/group, **○** (blue) male, ◇ (red) female) (C) Average food intake of male and female sedentary (SED) and ET *Slirp* KO or WT mice measured for 2 weeks after 4 weeks of running (n=3-8, **○** (blue) male, ◇ (red) female). (D-K) Blood glucose levels following a 4-hour fasting period before the glucose tolerance test (GTT). Glucose tolerance of male and female SED and ET *Slirp* KO or WT mice after 7-weeks of ET. iAUC of glycemic excursion in response to bolus of glucose 2 g kg^−1^ body weight (BW). Insulin response before (0 min) and following (20 min) the oral glucose challenge (n=6-11/group, **○** (blue) male, ◇ (red) female). Data are means ± SEM, including individual values where applicable. Geno, main effect of genotype; ET, main effect of exercise training; Acute exercise, main effect of acute exercise bout; Time X Geno ET, interaction between glucose bolus and genotype in the ET groups. *p<0.05, **p<0.01; (E) WT ET vs. KO ET, *p<0.05; WT SED vs. WT ET, #p<0.05, as per Unpaired Student’s t test (B), Two-way RM ANOVA with Šídák’s multiple comparisons test in ET groups (E, I), Two-way ANOVA (C, D, F, G, H, J, K).

The daily running distance recorded over 6 weeks of the 10-week study was 2.4 km/day on average for male mice of both genotypes, whereas it was 4.8 km/day for female exercise-trained (ET) WT mice and 3.0 km/day for female KO ET mice (Fig. 3B). Of note, the running distance observed in our study of 18-week old mice is shorter than those reported in other exercise training studies using ∼ 10-week-old mice^23,45,46^. Despite the higher food intake in ET mice (Fig. 3C), exercise training lowered body weight in both trained genotypes (Supplementary Fig. 2A). Moreover, irrespective of genotype, exercise training reduced fat mass/body weight ratio (Supplementary Fig. 2B) and increased lean mass/body weight ratio compared to sedentary (SED) groups (Supplementary Fig. 2C). Exercise training had no effect on gastrocnemius or TA muscle weight relative to tibia length in either sex (Supplementary Fig. 2D). However, we noted that female *Slirp* KO SED and both ET groups had elevated heart weight relative to WT SED (Supplementary Fig. 2D).

Exercise capacity increased similarly with exercise training, independently of genotype (+1.2-fold for WT ET, +1.3-fold for KO ET; Supplementary Fig. 2E) and sex (females shown in Supplementary Fig. 2G). Post-exercise blood lactate levels tended to be reduced (−1.7-fold; p=0.067) by ET in WT male mice (Supplementary Fig. 2F). The training-induced lowering of blood lactate post-exercise was absent in male and female *Slirp* KO mice (Supplementary Fig. 2F, H).

Exercise training is a powerful preventative treatment against many metabolic disorders^12^, partially due to its remarkable effects in improving glucose tolerance and insulin sensitivity documented in both animal models and humans^23,45,47^. Accordingly, male WT ET mice displayed reduced fasting blood glucose (Fig. 3D) and improved blood glucose control evidenced by a downshifted glucose response curve compared to WT SED mice. This effect was not observed in trained male *Slirp* KO mice (Fig. 3E). Thus, trained *Slirp* KO mice displayed similar glucose tolerance as both SED groups. The incremental area under the curve (iAUC) of blood glucose was similar between groups (Fig. 3F), suggesting that the lowered plasma glucose excursion during the GTT was driven by lower basal glucose levels. Yet, exercise training reduced the levels of plasma insulin in both WT and *Slirp* KO mice, suggesting that insulin sensitivity was increased by exercise training in both groups (Fig. 3G). There was no effect of exercise training or genotype on fasting glucose, glucose tolerance or insulin levels in female mice (Fig. 3H-K).

Our results demonstrate that SLIRP is largely dispensable for organismal adaptations to exercise training, yet, in male mice required for training-induced improvement in blood glucose regulation.

### Exercise training reverses *Slirp* KO-induced defects in muscle mitochondrial structure and function

Exercise training can counteract mitochondrial damage, arising from excessive accumulation of reactive oxygen species (ROS) or impaired assembly of OXPHOS complex following mtDNA mutations^11,13,48–50^. As we established that loss of SLIRP had marked negative implications for mitochondrial structure and respiration in skeletal muscle (Fig. 1), we asked whether exercise training could rescue these defects.

Skeletal muscle mitochondrial network structure was qualitatively investigated in TMRE+ stained FDB fibers by confocal microscopy. Remarkably, exercise training completely rescued the derangements in mitochondrial network structure and mitochondrial fragmentation observed in *Slirp* KO SED mice, quantitatively corroborated by restoration of the fragmentation index (Fig. 4A).

**Fig 4.**
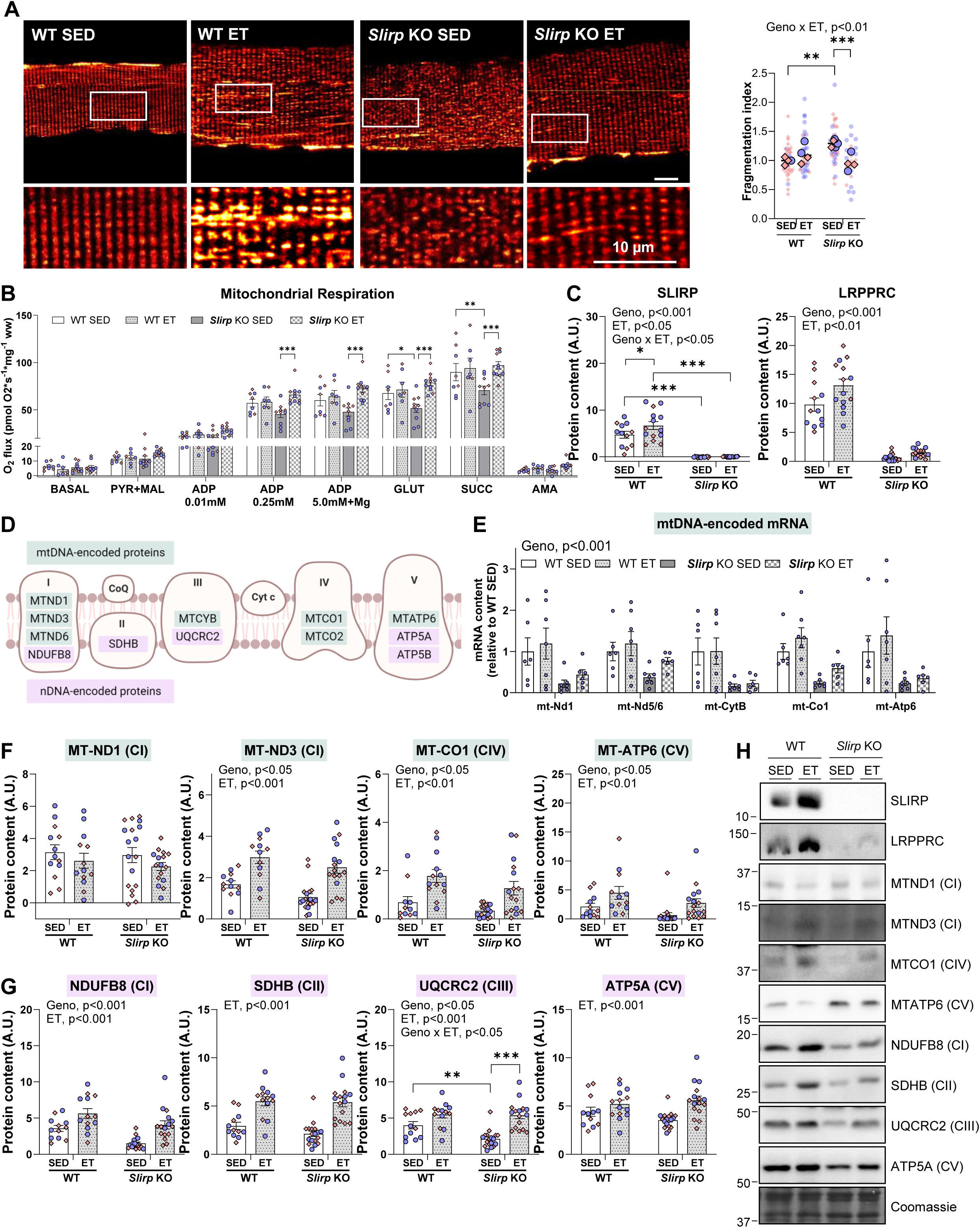
Exercise training reverses SLIRP-induced defects in muscle mitochondrial structure and respiration despite sustained reductions of mitochondrial transcripts. (A) Confocal microscopy of mitochondrial network structure using a mitochondrial membrane potential probe (TMRE+) in flexor digitorum brevis muscle fibers of sedentary (SED) and 10-week ET *Slirp* knockout (KO) mice and littermate controls (WT) (n=3-4, 7-10 fibers per mouse), and corresponding fragmentation index. SED data is also depicted in Fig. 1D. (B) Mitochondrial respiration measured by Oroboros respirometry system in gastrocnemius of SED and ET *Slirp* KO and WT (n=2-6/per group, **○** male, ◇ female). SED data is also depicted in Fig. 1H. (C) Western blot analysis of SLIRP and LRPPRC protein abundance in gastrocnemius of SED and 10-week ET *Slirp* KO and WT mice (n=6-11/group, **○** (blue) male, ◇ (red) female). (D) Schematic of mtDNA- and nuclear DNA-encoded oxidative phosphorylation (OXPHOS) proteins analysed. (E) RT-qPCR analysis of mitochondrial transcript levels in gastrocnemius of male SED and ET *Slirp* KO and WT mice (n=6-11/group). SED data is also depicted in Fig. 1J. (F-H) Western blot analysis of (F) mtDNA-encoded proteins, MT-ND1 (CI), MT-ND3 (CI), MT-CO1 (CIV), MT-ATP6 (CV), and (G) nDNA-encoded proteins NDUFB8 (CI), SDHB (CII), UQCRC2 (CIII) and ATP5A (CV), and (H) representative blots in gastrocnemius of SED and ET *Slirp* KO and WT mice (n=6-11/group, **○** (blue) male, ◇ (red) female). Data are means ± SEM, including individual values. Geno, main effect of genotype; ET, main effect of exercise training; Geno x ET, interaction between genotype and exercise training; Substrate, main effect of substrate. *p<0.05, **p<0.01, ***p<0.01 as per Two-way ANOVA with Šídák’s multiple comparisons test (A, B, C, F, G).

Concomitantly with improved mitochondrial structure, exercise training restored the respiratory flux in *Slirp* KO mice to the level of WT muscle (Fig. 4B). WT mice did not improve respiratory flux with exercise training, which was consistent with no effect of exercise training on fatty acid oxidation of contracting isolated WT soleus muscle (Supplementary Fig. 2I) and aligns with other studies^51^. Exercise training in *Slirp* KO mice improved fatty acid oxidation in contracting *Slirp* KO soleus muscles (Supplementary Fig. 2I), in alignment with improved respiratory flux in *Slirp* KO muscles by exercise training. Thus, exercise training counteracted the structural alterations in *Slirp* KO SED mice and had a positive impact on mitochondrial respiratory capacity, suggesting restored mitochondrial function.

Together, these results support that exercise training exploits the remarkable adaptive plasticity that mitochondria retain even in the event of mitochondrial network disruptions and impaired respiratory flux in skeletal muscle.

### Sustained reductions of mitochondrial transcript levels can be bypassed by exercise training

We next aimed to investigate the molecular underpinnings for the defects in mitochondrial structure and function in *Slirp* KO and their correction by exercise training. First, we verified that both SLIRP (+1.4-fold; Fig. 4C) and concomitantly LRPPRC (+1.4-fold; Fig. 4D) were upregulated by exercise training in the gastrocnemius muscle in WT, but not *Slirp* KO mice (Fig. 4C, D). These findings reinforce the co-stabilizing relationship of SLIRP and LRPPRC existing not only at baseline^17,18,20,52^ but also during exercise training conditions.

SLIRP, together with LRPPRC, has been shown to maintain mt-mRNA stability and aid mitoribosomal translation in non-muscle tissues^17–20,52^. Indeed, the marked downregulation of mt-mRNA transcripts in SED *Slirp* KO muscle, also shown in Fig. 1J, remained reduced in ET *Slirp* KO mice relative to WT ET mice (Fig. 4F). In contrast, mtDNA copy number, measured using quantitative PCR and primers for mt-*Nd1* (Supplementary Fig. 2J) or *mt-Nd5/mt-Nd6* (Supplementary Fig. 2K), were unaltered across groups or elevated only in *Slirp* KO SED mice, suggesting a compensatory response to mitigate mitochondrial dysfunction by increasing mtDNA copy number^53^. Thus, these findings suggests that SLIRP is crucial for mitochondrial transcript stability in skeletal muscle. We were intrigued to find that exercise training still restored protein content of mtDNA-encoded OXPHOS subunits, MT-ND3, MT-CO1 and MT-ATP6 (Fig. 4F). Thus, exercise training completely counteracted the sustained reduction in the mt-mRNA levels of these proteins in *Slirp* KO ET mice. The mild reduction of MT-ND3 protein, a subunit of Complex I, seen in *Slirp* KO muscle was similarly rescued by exercise training (Fig. 4F). Interestingly, although the nuclear-encoded Complex I subunit NDUFB8 was downregulated by 57% in the *Slirp* KO SED mice, exercise training markedly upregulated NDUFB8 protein content in *Slirp* KO mice, yet, below NDUFB8 protein content in WT ET mice (Fig. 4G). Other nuclear-encoded OXPHOS subunits, such as SDHB (Complex II), UQCRC2 (Complex III) and ATP5A (ATP synthase) were similarly upregulated by exercise training, independently of *Slirp* KO (Fig. 4G, representative blots shown in Fig. 4H). Citrate synthase (CS) activity correlates with mitochondrial content in human^54^ and mouse skeletal muscle^55^. CS activity, indicative of mitochondrial content, was increased similarly in *Slirp* KO and WT mice after exercise training (Supplementary Fig. 2L), suggesting that improvements in mitochondrial content following exercise training are independent of SLIRP.

These results underline a substantial skeletal muscle mitochondrial plasticity, which is retained despite a 60-80% reduction in mt-mRNA pools. These findings point towards unidentified mechanisms by which ET improves translational efficiency to dictate final protein content.

### Exercise training increases mitochondrial protein translation capacity by elevating mitoribosome mass

We next sought to determine the potential mechanism by which sustained reductions of mitochondrial transcript levels can be bypassed by exercise training. Enhanced mitochondrial protein translation mediates exercise training adaptations in muscle mitochondrial function in both young and elderly individuals^39^. Thus, we assessed pathways associated with protein translation in mitochondria, facilitated by the resident mitoribomes. We observed training-induced increases in 12S ribosomal RNA (12S rRNA) and MRPL11 protein content (Fig. 5A, B), essential components of mitoribosome translation^56,57^. MRPL11 protein content was 36% more elevated in response to exercise training in *Slirp* KO ET than WT ET mice, mainly in male mice (Fig. 5B). Conversely, exercise training did not induce changes in cytosolic ribosomal protein S6 protein content (rpS6, Fig. 5C; representative blots in Fig. 5D), indicating a compartmentalized ribosomal response to exercise training.

**Fig 5.**
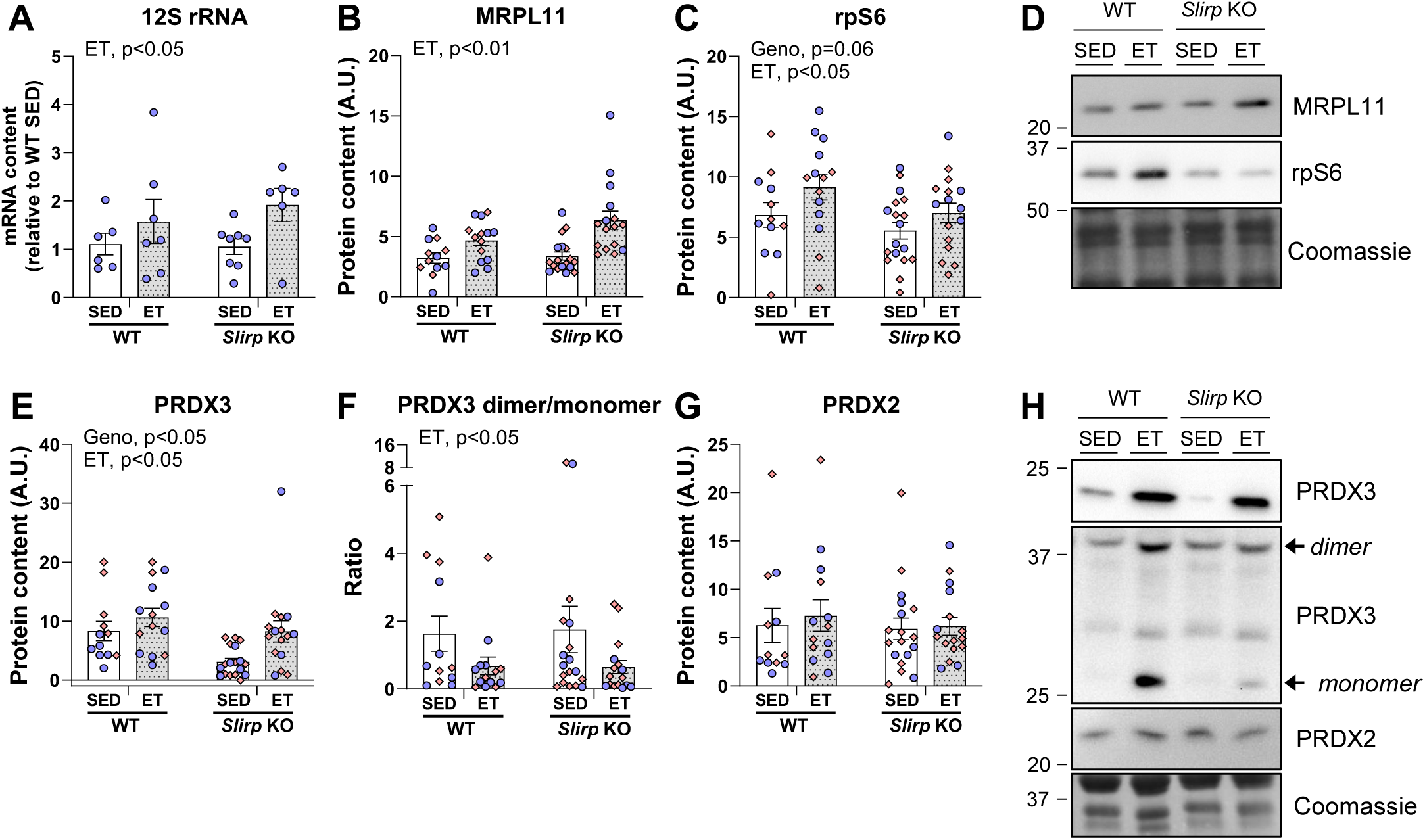
Exercise training increases mitochondrial protein translation capacity by elevating mitoribosome mass. (A-H) RT-qPCR analysis of 12S rRNA (A) and Western blot analysis of MRPL11 (B), rpS6 (C), PRDX3 (E), PRDX3 dimer/monomer ratio (F), PRDX2 (G), and representative blots in gastrocnemius of sedentary (SED) and exercise trained (ET) *Slirp* knockout (KO) and control littermate (WT) mice (n=6-11/group, **○** (blue) male, ◇ (red) female). Data are means ± SEM, including individual values. Geno, main effect of genotype; ET, main effect of exercise training; *p<0.05, **p<0.01, as per Two-way ANOVA (A-C, E-G).

Given the predominant impact of *Slirp* KO on mitochondria, training-induced adaptive changes might also manifest directly within the affected cellular compartment. Here, we investigated a mitochondrial quality control system involving a mitochondrial scavenger enzyme, peroxiredoxin 3 (PRDX3). PRDX3 plays a role in balancing redox environment in mitochondria^58,59^, possibly contributing to the preservation of factors critical for translation during conditions of mitochondrial dysfunction and exercise training. Interestingly, PRDX3 protein content was 60% lower in *Slirp* KO SED compared to WT SED mice, indicative of a lower mitochondrial oxidative stress defense capacity in *Slirp* KO muscle. This was restored by exercise training, bringing PRDX3 up to levels observed in trained WT mice (Fig. 5E). We further noted that ET lowered the PRDX3 dimer to monomer ratio in both genotypes, indicative of a lower basal mitochondrial-derived peroxide accumulation likely due to an enhancement of scavenger activity (Fig. 5F). The cytosolic peroxidoxin 2 (PRDX2) was unaffected by genotype and exercise training (Fig. 5G, representative blots in Fig. 5H), further pointing towards highly mitochondria-selective effects of exercise training to circumvent SLIRP deficiency.

Collectively, our findings indicate a complex interplay of spatially distinct molecular muscle adaptations in response to *Slirp* KO and ET. Our findings suggest that exercise training induces mechanisms to increase mitochondrial protein translation capacity. Mechanistically, exercise training elevated mitoribosome mass and restored oxidative stress defense systems to powerfully circumvent the *Slirp* KO-induced depletion of mtDNA-encoded transcript levels.

### Elevation of muscle SLIRP and LRPPRC protein content in response to ET is conserved in human skeletal muscle

Our intriguing results from mouse and fly models prompted us to test whether our results provide translational value for humans in health and disease. SLIRP and LRPPRC protein content were determined in human skeletal muscle in four independent exercise cohorts employing different exercise modalities in conjunction with or without aging or diabetes and in males and females. Participant characteristics and study designs for the different ET modalities have been published previously for all the clinical studies^60–63^.

Skeletal muscle SLIRP and LRPPRC protein abundances were 70% increased following 14-weeks of controlled and supervised aerobic and strength exercise training intervention in healthy young women^60^ (Fig. 6A). Also, a 6-week high-intensity interval training (HIIT) intervention elevated muscle protein content of SLIRP (+80%, Fig. 6B) and LRPPRC (+50%, Fig. 6B) in healthy young men^63^. Thus, the exercise training response of SLIRP was highly consistent, independent of sex, and responsive to various exercise modalities. The 12-week progressive resistance exercise training^61^ increased SLIRP and LRPPRC in the entire group of young and old individuals but had no significant effect on LRPPRC abundance in either young or elderly subjects separately (Fig. 6C). The mtDNA-encoded transcript levels of *MT-ND6*, *MT-CYB*, *MT-CO1* and *MT-ATP6* were comparable across all groups (Fig. 6D), indicating that mitochondrial gene expression was not significantly affected by either aging or the training regimen in this study.

**Fig 6.**
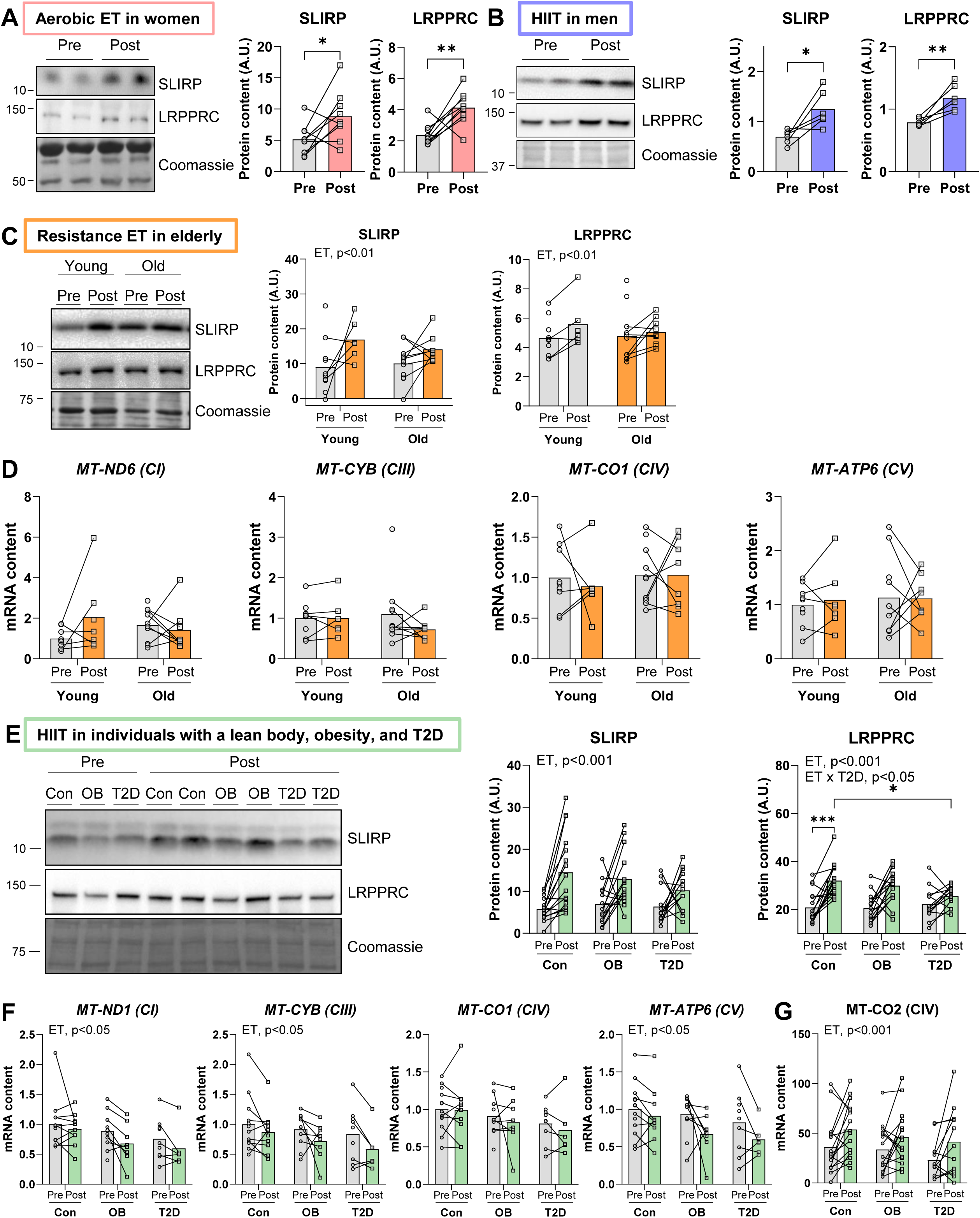
In human skeletal muscle SLIRP and LRPPRC protein content are increased by exercise training. (A) Western blot analysis of SLIRP and LRPPRC protein in vastus lateralis of healthy young women (n=9/group, (**○** Pre ET, □ Post ET)) after a 14-week controlled aerobic and strength exercise training (ET) intervention^60^.(B) Immunoblotting of SLIRP and LRPPRC protein in vastus lateralis of healthy young men (n=6/group, (**○** Pre ET, □ Post ET)) after a 6-week high intensity interval training^63^.(C, D) Immunoblotting of SLIRP and LRPPRC protein (C) and RT-qPCR analysis of mitochondrial transcript levels (D) in vastus lateralis of male young and older individuals after 12-week progressive resistance training^61^ (n=6-11/group, (**○** Pre ET, □ Post ET)).(E-G) Immunoblotting of SLIRP, LRPPRC (E), and MT-CO2 (G) protein and RT-qPCR analysis of mitochondrial transcript levels (F) in vastus lateralis of male individuals, who were lean, obese or type 2 diabetic after high-intensity interval training^62^ (n=13-16/group**○** pre ET, □ post ET). Con, control; OB, obese; T2D, Type 2 diabetes.Data presented as individual before-and-after values. ET, main effect of exercise training, ET x T2D, Interaction between exercise training and presence of type 2 diabetes in Con and T2D group; *p<0.05,**p<0.01, ***p<0.001, as per paired Student’s t test (A-C), Mixed-effects model (D, F), Two-way RM ANOVA with Šídák’s multiple comparisons test (E, G).

Exercise training-mediated improvements in mitochondrial content and function may be associated with improved glucose metabolism in individuals with obesity and T2D ^64,65^, although this association is not always clear^66^. The training response of SLIRP and LRPPRC to HIIT^62^ was conserved in individuals who were lean (+150% and +54%) or obese (+85% and +45%, Fig. 6E). However, the exercise response for LRPPRC, but not SLIRP, was blunted in patients with T2D in comparison to lean individuals (Fig. 6E), suggesting impaired sensitivity of muscle to some mitochondrial adaptations to ET in T2D. At the mRNA level, mt-DNA encoded transcripts of *MT-ND1*, *MT-CYB*, *MT-CO1* and *MT-ATP6* were reduced with HIIT across all groups (Fig. 6F). Yet, protein content of MT-CO2 (Fig. 6G, representative blot in Supplementary Fig. 3A) and other nuclear-encoded OXPHOS proteins (Supplementary Fig. 3A) were significantly upregulated with HIIT. In line with previous findings^39^, the upregulation in OXPHOS protein abundance occurred despite lower levels of mRNA, and demonstrate a disassociation between the transcriptome and proteome^67^.

Together, we show that SLIRP and LRPPRC are robustly upregulated in skeletal muscle across various exercise training modalities in men and women, lean and obese. However, the exercise training response of LRPPRC was blunted in patients with T2D. This suggests that T2D is associated with decreased sensitivity to some of the adaptive mitochondrial responses normally elicited by exercise training, possibly influenced by exercise intensity or volume. Thus, our clinical data in humans provide strong translational value of the SLIRP-deficient fly and mouse models and point towards a conserved requirement of SLIRP for skeletal muscle health, and potentially the SLIRP-independent mechanisms that compensate during exercise training.

## Discussion

Our investigation led to six key findings that bridge substantial knowledge gaps concerning the role of mitochondrial posttranscriptional mechanisms on skeletal muscle biology and their contributions to exercise training adaptations. Our findings not only unravel a delicate interplay between the mt-mRNA-stabilizing protein SLIRP, and the integrity of skeletal muscle mitochondria structure and performance, but also underscore the potential of exercise training in promoting selective mitochondrial translation capacity to circumvent these mechanisms.

First, lack of *Slirp* led to mitochondrial network disruption, mitochondrial fragmentation, and reduced respiratory capacity in skeletal muscle in mice. Second, SLIRP deficiency impaired muscle functionality, and reduced lifespan in flies. Third, SLIRP was identified as an exercise training-responsive downstream target of PGC1-α. Fourth, SLIRP was needed for improving blood glucose regulation after exercise training in male mice. Fifth, exercise training restored the mitochondrial defects elicited by the lack of SLIRP on mitochondrial structure, respiration, and mtDNA-encoded OXPHOS, via mechanisms likely involving the upregulation of mitoribosome content and enhanced antioxidant defense. Finally, we established a translational foundation for our findings by substantiating the conservation of SLIRP’s response to exercise training in four independent human cohorts integrating different exercise modalities, sex, health, and age conditions.

Our first finding establishes SLIRP as an hitherto unrecognized player in regulating mitochondria content, structure, and respiration in skeletal muscle. Our results are partially consistent with reports in heart, liver, and kidney *Slirp* KO^17,18,52^ showing that SLIRP forms a complex with LRPPRC and is crucial for the maintenance of the mt-mRNA reservoir. Accordingly, our observation of mitochondria with abnormal cristae in *Slirp* KO muscle aligns with transmission electron micrographs of *Lrpprc* KO heart tissue^19^. However, despite the stark similarities, mitochondrial respiration was compromised in *Slirp* KO skeletal muscle, but not in heart or liver^18^, underscoring SLIRP’s functional importance within skeletal muscle relative to other tissues. In alignment, previous research further showed that *Slirp* KO skeletal muscle fibers have reduced sarcoplasmic reticulum Ca^2+^ storage capacity^68^. Thus, our present findings underscore the importance of SLIRP across various tissue types, with emphasis on its importance to skeletal muscle mitochondrial morphology and respiration, and overall mitochondrial health.

Our second finding that SLIRP depletion compromised climbing ability, impaired starvation tolerance, and reduced lifespan in flies, illustrates the detrimental functional consequences of lacking SLIRP in the muscle tissue. The effects of mitochondrial proteins on modulating life span are equivocal. Some report that the depletion of specific OXPHOS genes reduces life span, whereas others report no effect or an increase on life span in model organisms^69–71^. The “Mitochondrial Threshold Effect Theory”^72^ posits that mild mitochondrial dysfunction allows normal physiology in model organisms until a threshold is reached. Beyond that, severe mitochondrial dysfunction can lead to premature aging, developmental arrest or even death^73–75^. In support, heterozygous depletion of *Mclk1*, which encodes an enzyme necessary for biosynthesis of the electron carrier coenzyme Q of OXPHOS, even increased lifespan in mice, but homozygous depletion was embryonically lethal^75^. Our findings support an important role for SLIRP and mitochondrial function in maintaining normal lifespan because we observed reduced survival in muscle-specific SLIRP KD flies.

Our third finding was that PGC-1α1 regulated SLIRP protein, providing evidence for a new training-responsive PGC-1α target in skeletal muscle. Interestingly, training-induced adaptations in mitochondrial proteins can occur independently of PGC-1a1 as the main regulator^76^. In agreement with that, our fifth finding was that the training-induced improvements in mitochondrial function could bypass SLIRP deletion. These findings agree with other studies showing that exercise training can rescue mitochondrial dysfunction and elicit improvements in glucose homeostasis in mice and humans^13,48,76,77^. Interestingly, SLIRP was required for the training-induced improvements in blood glucose regulation in trained male *Slirp* KO mice. While not explored in our study, this could be due to decreased intramyocellular insulin signalling^78^, reduced insulin-independent glucose uptake^14^, capillarization, and/or blood flow^79^. Female mice did not improve their glucose tolerance in response to exercise training. Sexual dimorphism in response to metabolic challenges, including exercise training, has been frequently reported in mouse models^80,81^. Yet, in both sexes, SLIRP is largely dispensable for exercise training-induced mitochondrial and metabolic adaptations in skeletal muscle, even though SLIRP is crucial for mitochondrial transcript stability in the basal state.

We were intrigued by the observation that exercise training could circumvent even 60-80% loss in mitochondrial-mRNA pools to upregulate mtDNA-encoded OXPHOS proteins. In other words, the beneficial effect of exercise training on mitochondrial oxidative phosphorylation and ATP provision is independent of a correction of mt-mRNA transcript levels and LRPPRC/SLIRP protein content. These findings give rise to the possibility that mt-mRNAs are produced in excess *in vivo*^18^. Excess mt-mRNA may allow for rapid engagement of mitochondrial protein synthesis in the event of sudden changes in energy demand. In response to exercise training, the mitoribosomal components, MRPL11 and *12S rRNA* were upregulated, indicating increased capacity for mitochondrial protein synthesis. These adaptations to exercise training may help sustain protein translation, even in the presence of low mt-mRNA abundance. Exercise training also upregulated PRDX3, linked to peroxide scavenging. While MRPL11, *12S rRNA* and PRDX3 may not directly control translation efficiency, their roles in the mitoribosomal makeup and improved redox state can influence the overall cellular environment where protein translation occurs. Albeit limited by not directly measuring mitochondrial translational rate, this working hypothesis is supported by findings in human skeletal muscle showing that mitochondrial protein translation is at the core of exercise training-induced benefits^39^.

Finally, our results likely are relevant to humans, as SLIRP and LRPPRC were consistently upregulated in human skeletal muscle in response to multiple exercise training modalities in both sexes. The effect of resistance exercise training was blunted for LRPPRC in patients with T2D. Interestingly, the age-related decline in muscle mitochondrial protein synthesis^82^ can be reversed by exercise training in elderly^39^, corroborating our findings that ET bypasses SLIRP to increase mitoribosomal translation. Importantly, our findings demonstrate that the synthesis of mtDNA-encoded OXPHOS proteins is not limited by transcript abundance in skeletal muscle – a feature that seems to be conserved as a fundamental mechanism^26,39^.

Taken together, our findings not only unravel a new mechanism for the regulation of basal mitochondrial function in skeletal muscle, but also highlight an incredible exercise training –stimulated plasticity of mitochondria in skeletal muscle facing mitochondrial defects. Our findings underscore ET as a therapeutic intervention to combat mitochondrial dysfunction in genetic and lifestyle-induced muscle pathologies, including T2D.

## Conclusions

In this work, we uncover a conserved role for mitochondrial transcript stability via SLIRP in basal mitochondrial structure and respiratory capacity in skeletal muscle. We identify SLIRP as a novel exercise-responsive downstream target of PGC-1α that is conserved in human skeletal muscle. Because the defects in *Slirp* KO muscle could be reversed by ET, these findings add to the clinical relevance and add impetus to the important health-promoting role of ET in pathologies characterized by mitochondrial defects.

## Methods

### Fly maintenance

In standard conditions, the flies were maintained on a SYA medium containing 0.5% (w/v) agar, 2.4% (v/v) nipagin, 0.7% (v/v) proprionic acid, 10% (w/v) dry baker’s yeast and 5% (w/v) sucrose. The experiments took place at +25°C, 65% humidity under a 12h:12h light:dark cycle. RNAi lines for *SLIRP1* (51019 GD), *SLIRP2* (23675 GD), and the GD library control line (w1118, 60000 GD) were obtained from Vienna Drosophila Resource Center. Mef2-GAL4 line was obtained from Bloomington Stock Center to generate muscle-specific *SLIRP* KD flies.

### Negative geotaxis (climbing) assay

*Drosophila* has a natural tendency to climb upwards, also known as negative geotaxis. The negative geotaxis, or climbing, assay was performed as previously described^83^. Briefly, 20 male *Drosophila* flies aged 5-6 days were transferred into empty vials (25 x 95mm), and another empty vial was taped on top to create a 25 x 190mm cylinder. Six cylinders were set side by side into an apparatus. The apparatus was tapped down 5 times to make all the flies fall to the bottom of the cylinder. The climbing assay was measured in technical triplicate for each biological replicate, with 90 seconds allowed for each climb. The assay was recorded with a camera and videos were analysed to determine the time for half of the flies in the vial to reach halfway point (95mm) in the cylinder.

### Starvation assay using the Drosophila Activity Monitoring System (DAMS)

Four-day old male flies were individually housed in monitor tubes with an outside diameter of 5mm (PPT5×65, Trikinetics, Waltham, MA). For starvation analysis, the tubes contained 1% agar in water. 32 tubes per monitor (DAM2 Drosophila Activity Monitor, Trikinetics, Waltham, MA) were set up, one monitor per genotype. For the starvation analysis, flies were kept in the DAM system until the last fly had died.

### Lifespan assay

For lifespan assays, flies were mated at a controlled density of 25 female and 10 male flies, respectively. One-day old male flies were collected into vials (10 male flies per vial, with 10 vials per genotype). The flies were kept on a SYA diet in Drosoflippers (http://www.drosoflipper.com/). Flies were flipped onto fresh food every second day and deaths were scored during the transfer.

### Mice

#### PGC-1α1 transgenic mouse (PGC-1α1 OE)

The generation of the model, quality, and specificity of the overexpression has been described previously^35^, of which we analyzed quadriceps muscle samples.

#### PGC-1α4 transgenic mouse (PGC-1α4 OE)

The generation of the model, quality, and specificity of the overexpression has been described previously^33^, of which we analyzed gastrocnemius muscle samples.

#### Skeletal muscle-specific PGC-1α knockout (PGC-1α mKO)

The generation of the model, quality, and specificity of the KO has been described previously^44^. Male muscle-specific PGC-1α MKO mice and littermate control mice homozygous for loxP inserts (lox/lox) were generated by crossbreeding myogenin-Cre mice with loxP flanked-Pgc-1α mice.

For the acute exercise bout, Tibialis anterior muscles of PGC-1α MKO mice and littermate control mice were collected 3 hours after one bout of equal distance treadmill running (1.4 km) at 10° incline and 60% of their individual maximal running speed achieved by a graded treadmill running test^84^.

For the ET intervention, mice were single-housed with or without access to in-cage running wheels from 8 to 20 weeks old. Running distance and duration were monitored by a regular cycle computer. PGC-1α MKO mice tended to run less than lox/lox, thus running wheels of lox/lox mice were occasionally blocked to ensure equal running distance. On average, mice ran 25 km/week. The running wheels were blocked for all mice 24 hours prior to euthanization and harvest of quadriceps muscle in the morning in the fed state.

#### AAV6 Vector construction and preparation

The AAV6 vector for SLIRP overexpression and ultra-purified eGFP control AAV6 virus were manufactured by VectorBuilder Inc. (Shenandoah, Texas, USA; AAV6SP(VB190219-1010jvy)-C).

#### Transfection of tibialis anterior muscle using rAAV6 vector

Viral particles were diluted in Gelofusine (B. Braun, Germany) to a dosage of 5 × 10^9^ vector genomes at a volume of 30 uL per injection. For muscle-specific delivery of rAAV6 vectors, 12-week-old mice were placed under general anaesthesia (2% isofluorane in O_2_) and mice were injected intramuscularly with rAAV6:SLIRP in the TA muscle of one leg and control rAAV6:eGFP (empty vector) in the contralateral leg. Muscles were harvested 4 weeks after rAAV6 administration.

#### Time course of acute exercise

Twelve-week old female C57BL/6J mice (n=8 for each time point) were subjected to an acute exercise bout (1h running, 60% of maximal running intensity, 15° incline). Quadriceps muscle was harvested from rested and exercised mice immediately after, 2h, 6h, and 24 h after the acute exercise bout.

#### SLIRP knockout mouse (*Slirp* KO mice)

The generation of the model, quality, and specificity of the KO has been described previously^18^. Sperm of homozygous *Slirp* KO mice kindly provided by Nils-Göran Larsson and re-derived offspring was backcrossed to C57BL/6N background in our own animal facilities.

All mice were maintained under a 12:12 h light/dark photocycle at 22 ± 2°C with nesting material. The female mice were group-housed (except during the ET intervention), whereas the male mice were single-housed. All mice received a rodent chow diet (Altromin no. 1324; Chr. Pedersen, Denmark) and water *ad libitum*.

Mouse genotyping of *Slirp* KO and WT mice was performed as previously described ^85^ using qPCR on DNA from ear punches with the following primers: WT; 5’-AGAAGGGAAT CCACAGGATA GGACA-3’ and 5’-GCTTTATTCC TAGTGCTGGC CTTGTT-3’, KO; 5’-AGAAGGGAAT CCACAGGATA GGACA-3’ and 5’-CGCCGTATAA TGTATGCTAT ACGAAGTT-3’.

#### 10-week ET intervention in *Slirp* KO mice

For 10-week voluntary wheel-running exercise training interventions, 16–18-week-old ***Slirp*** KO and littermate mice were randomized into test groups with or without access to in-cage running wheels (Tecniplast activity cage, wheel diameter: 23 cm; Tecniplast, Buguggiate VA, Italy). The experimental design is schematically illustrated in Fig. 3A. Running distance was monitored by a regular cycle computer prior to the metabolic tests for 6 weeks. Running wheels were locked 12 h prior to terminal procedures to avoid effects of acute exercise bouts.

#### Body composition

Total, fat, and lean body mass was measured by nuclear magnetic resonance using an EchoMRI™ (USA).

#### Glucose tolerance test (GTT) and plasma insulin analysis

We subjected the *Slirp* KO and WT mice to a GTT at 7 weeks of the training intervention study (as illustrated in Fig. 3A). The GTT was executed after a 5 h fasting period (7:00 a.m.–12:00 p.m.). Resting blood samples were taken from the tail 30 min prior to the intraperitoneal injection of d-mono-glucose (2 g/kg body weight). Tail blood glucose was measured after 0, 20, 40, 60, and 90 min of injection.

To measure glucose-stimulated plasma insulin concentration at time-point (min) 0 and 20, tail vein blood samples were collected in a capillary-tube (50 µl), centrifuged at 14,200 g for 5 min at 4 °C, plasma collected and stored at −80°C. Insulin concentration was determined in duplicates using the Ultra-Sensitive Mouse Insulin ELISA Kit (#80-INSTRU-E10; ALPCO Diagnostics) to the manufacturer’s instructions. The incremental area under the curve (iAUC) from the basal blood glucose concentration was determined using the trapezoid rule.

#### Exercise capacity tests

*Slirp* KO and WT mice were acclimatized to the treadmill on 3 consecutive days by running at a speed of 0.16 m/s and incline of 10° for 5 min the first day and 10 min the following days. Prior to the running test and with a 1h delay, 2 blood samples were drawn from the tail before the test for pre-exercise blood glucose and blood lactate measurements. Afterwards, the mice ran at 0.16 m/s for 5 minutes followed by a gradual increase in speed every minute with 0.02 m/s until exhaustion. Once the mouse reached its maximal running capacity, 2 blood samples were immediately drawn from the tail for post-exercise blood glucose and blood lactate measurements. The test was stopped when the mouse failed to keep up with the treadmill despite motivational efforts by the researcher.

#### TMRE staining in live fibers

Flexor digitorum brevis muscles (FDB) were dissected and then incubated for 2 hours in serum-free α-MEM (22571-020, Gibco) containing 1.9 mg/ml collagenase type 1 from clostridium histolyticum (C0130, Sigma-Aldrich) and 1 mg/ml bovine serum albumin (9048-46-8, Sigma-Aldrich) at 37° C on a rotator. After collagenase treatment, muscles were incubated in α-MEM containing 10% fetal bovine serum (26050-70, Gibco) and subjected to mechanical dissociation using fire-polished Pasteur pipettes. Single muscle fibers were seeded in 35×14mm glass-bottom microwell dishes (P35G-1.5-14-C, MatTek Corporation) coated with 4 uL of Engelbreth-Holm-Swarm murine sarcoma ECM gel (E1270, Merck). Muscle fibers were kept in α-MEM containing 5% fetal bovine serum in a cell incubator (37°C, 5% CO2) for at least 16 h prior to the experiments.

For determination of mitochondrial membrane potential (ΔΨmitochondrial) and mitochondrial network morphology, the fibers were incubated in 20nM tetramethylrhodamine and ethyl ester (TMRE+, Life Technologies) dissolved in Krebs Ringer buffer (145 mM NaCl, 5 mM KCl, 1 mM CaCl_2_, 1 mM MgCl_2_, 5.6 mM glucose, 20 mM HEPES, pH 7.4) for 30 minutes before imaging. Confocal images were collected using a C-Apochromat ×40, 1.2 NA water immersion objective lens on an LSM 980 confocal microscope (Zeiss) driven Zeiss Zen Blue 3.

Mitochondrial network analysis was performed semi-automatically in ImageJ (National Institute of Health, USA). At least two nucleus-free regions per fiber were analyzed blindly. Before segmentation, the background was subtracted, and the pixels were averaged using a value of mean=2. The images were segmented, and particle analyses revealed the relative area of the TMRE+ signal compared to the total area (% mitochondrial area). The fragmentation index was calculated as the number of objects in relation to the total area covered by the dye. The ImageJ ‘Red Hot’ lookup table was used to visualize the images.

#### Transmission electron microscopy analysis

A small longitudinal section (<2 mm) of the red portion of the gastrocnemius muscle tissue was fixed by immersion 2% glutaraldehyde in 0.05M Phosphatebuffer, pH 7.4 and stored at 4°C. The samples were rinsed four times in 0.1 M sodium cacodylate buffer, pH 7.4, and post-fixed with 1% osmium tetroxide (OsO_4_) and 1.5% potassium ferrocyanide [K_4_Fe(CN)_6_] in 0.1 M sodium cacodylate buffer, pH 7.4 for 90 min at 4°C. The samples were then rinsed twice and dehydrated through a graded mixture of alcohol at 4–20°C, infiltrated with graded mixtures of propylene oxide and Epon at 20°C, and embedded in 100% Epon at 30°C, as previously described (Nielsen et al., 2011). The samples were cut in 60 nm longitudinal sections using a Leica Ultracut UCT ultramicrotome. The sections were contrasted with uranyl acetate and lead citrate, and subsequently examined and image recorded in a CM100 TEM (Philips, Eindhoven, The Netherlands) equipped with a Veleta camera and the iTem software package (Olympus, Hamburg, Germany) with a resolution of 2048 x 2048 pixels.

A mean of eight fibers per sample (7-9) was included from the sectioned samples, and from each fiber, 24 images were obtained at x13,500 magnification. The imaging was performed in a randomized systematic order including 12 images from the subsarcolemmal (SS) region, and 6 from both the superficial and central region of the intermyofibrillar (IMF) space. The Z-disc width was measured once on all IMF images and the mean from all fibers within one sample was calculated to determine the fiber type. The categorization of fiber types was based on previous reports from observations^43^. The images were analyzed by two genotype-blinded investigators and all further analyses were performed by the same blinded investigator. The quantification of mitochondrial morphology was done using the Radius EM Imaging Software (Emsis GmbH, Radius 2.0).

Several measurements were included in this study: 1) Mitochondrial number: Total number of mitochondrial profiles (count). 2) Mitochondrial volume fraction: IMF mitochondrial volume is annotated as percentage of mitochondria covering the IMF space. 3) Damaged mitochondria: Mitochondria were categorized as damaged when they were swollen, vacuolated, or empty.

#### Mitochondrial respiration in gastrocnemius muscle

In a subset of mice, mitochondrial respiratory capacity was measured in permeabilized gastrocnemius skeletal muscle fibers as previously described^86^. In brief, gastrocnemius muscle was rinsed from fat and connective tissue and separated into small fiber bundles. Fiber bundles were permeabilized with saponin (50 μg/ml) in BIOPS buffer for 30 min, followed by a 20 min wash in MiR05 buffer on ice. Mitochondrial respiration was measured in duplicate under hyperoxic conditions at 37°C using high resolution respirometry (Oxygraph-2k, Oroboros Instruments, Innsbruck, Austria). The following protocol was applied: Leak respiration was assessed by addition of malate (5 mM) and pyruvate (5 mM), followed by adding three different concentrations of ADP (0.01 mM, 0.25 mM and 5 mM; for the final concentration of ADP, 3 mM of magnesium (Mg) was added as well) to measure complex I linked respiratory capacity. 10 mM Glutamate was added to measure maximal complex I linked respiratory capacity followed by 10 mM succinate to measure complex I+II linked respiratory capacity. Finally, 5 μM Antimycin A was added to inhibit complex III in the electron transport chain. Pooled data for both sexes are shown as no sex-specific differences were detected.

#### Contraction-stimulated fatty acid oxidation in isolated soleus muscles

Contraction-stimulated exogenous palmitate oxidation in isolated soleus muscle from *Slirp* KO and WT mice was measured as previously described ^87^. In brief, excised soleus muscles from mice anesthetized with pentobarbital were mounted at resting tension (∼5 mN) in 15 ml vertical incubation chambers with a force transducer (Radnoti, Monrovia, CA) containing 30°C carbonated (95% O_2_ and 5% CO_2_) Krebs-Henseleit Ringer buffer (KRB), pH=7.4, supplemented with 5 mM glucose, 2% fatty acid-free BSA, and 0.5 mM palmitate. After ∼20 min of pre-incubation, the incubation buffer was refreshed with KRB additionally containing [1–14C]-palmitate (0.0044 MBq/ml; Amersham BioSciences, Buckinghamshire, U.K.). To seal the incubation chambers, mineral oil (Cat. No. M5904, Sigma– Aldrich) was added on top. Exogenous palmitate oxidation was measured simultaneously at rest and during 25 min contractions (18 trains/min, 0.6 s pulses, 30 Hz, 60 V). After incubation, incubation buffer and muscles were collected to determine the rate of palmitate oxidation as previously described^86,88,89^. Palmitate oxidation was determined as CO_2_ production (complete FA oxidation) and acid-soluble metabolites (representing incomplete FA oxidation). As no difference was observed in complete and incomplete FA oxidation between genotypes, palmitate oxidation is presented as a sum of these two forms.

#### Lysate preparation and immunoblotting

Lysate preparation and immunoblotting of vastus lateralis muscles samples of HIIT in young, healthy men were performed as described by Hostrup et al.^90^, whereas lysate preparation and immunoblotting of vastus lateralis muscles of HIIT in individuals, who were lean, obese or T2D were performed as described by Kruse et al^91^.

To preserve and assess PRDX3 dimer to monomer ratio, freshly harvested mouse gastrocnemius muscle tissue was incubated for 10 min in ice-cold 100 mM N-Ethylmaleimide (NEM) diluted in PBS. NEM was subsequently aspirated and the tissue homogenized for 1 min at 30 Hz using a TissueLyser II bead mill (QIAGEN, USA) in ice-cold homogenization buffer [10% glycerol, 1% NP-40, 20 mM sodium pyrophosphate, 150 mM NaCl, 50 mM Hepes (pH 7.5), 20 mM β-glycerophosphate, 10 mM NaF, 2 mM phenylmethylsulfonyl fluoride, 1 mM EDTA (pH 8.0), 1 mM EGTA (pH 8.0), 2 mM Na_3_VO_4_, leupeptin (10 μg ml^−1^), aprotinin (10 μg ml^−1^), and 3 mM benzamidine]. Lysate preparation and immunoblotting of all remaining mouse tissues and human vastus lateralis muscle samples of aerobic and strength ET in women^60^, and resistance training in the young and elderly cohorts^92^ were performed as described in^23^. Immunoblotting of the NEM-treated samples was performed under non-denaturing conditions. The primary antibodies used are listed in Supplementary Table 2.

Coomassie Brilliant Blue staining was used as a control to assess total protein loading and transfer efficiency^93^ by quantifying the whole lane and for each sample set, a representative membrane from the immunoblotting is shown. The same Coomassie brilliant blue staining is presented for proteins analyzed when derived from the same sample set. Band densitometry was carried out using Image Lab (version 4.0). For each set of samples, a standard curve was loaded to ensure quantification within the linear range for each protein probed for. For immunoblotting in Fig. 5, pooled data for both sexes are shown as no sex-specific differences were detected, unless specified.

#### RNA extraction and RT–qPCR

Fig. 1J, 4F: RNA was extracted from ∼20 mg of pulverized whole mouse gastrocnemius muscle using TRIzol™ reagent (Invitrogen) following the manufacturer’s instructions (tissue homogenization was performed using the MP Bio lysis system) and treated with the TURBO DNA-free™ Kit (Ambion) to remove contaminating DNA. For RT–qPCR expression analysis, cDNA was reversed transcribed from 0.85 μg total RNA using the High-Capacity cDNA Reverse Transcription Kit (Applied Biosystems) in the presence of RNase Block (Agilent). The qPCR was performed in a QuantStudio 6 Flex Real-Time PCR System (Life Technologies), using TaqMan™ Universal Master Mix II, with UNG (Applied Biosystems) to quantify mitochondrial transcripts (mitochondrial-rRNAs and mt-mRNAs). The gene expression levels were determined using the ΔΔCt method, comparing the Ct values of mitochondrial transcripts to that of the beta-actin reference gene for normalization.

Fig. 2A, C, D: Total RNA was extracted from pulverized quadriceps muscle using an adapted guanidinium thiocyanate-phenol-chloroform extraction method. Reverse transcription to cDNA was performed as previously described^46^. Real-time qPCR was performed in triplicate with QuantStudio 7 Flex Real-Time PCR System (Applied Biosystems, Waltham, MA). Cycle threshold (Ct) was converted to a relative amount using a standard curve derived from a serial dilution of a representative pooled samples run together with the samples of interest. Beta-actin or Hprt mRNA was used for normalization of target mRNA levels.

Fig. 6J-M: Total RNA extraction from vastus lateralis muscle and reverse transcription to cDNA was performed as previously described^61^. The qPCR was performed in a QuantStudio 6 Flex Real-Time PCR System (Life Technologies), using TaqMan™ Universal Master Mix II (Applied Biosystems) to quantify mt-mRNAs. The gene expression levels were determined using the ΔΔCt method, comparing the Ct values of mitochondrial transcripts to that of the 18S rRNA reference gene for normalization.

Fig. 6Q-T: Total RNA was extracted from skeletal muscle biopsies using TRI Reagent (Sigma-Aldrich) following the manufacturer’s instructions. cDNA was reversed transcribed using the High-Capacity cDNA Reverse Transcription Kit (Applied Biosystems), while the qPCR was performed on an Aria Mx (Agilent) using a TaqMan™ Universal Master Mix II (Applied Biosystems). The gene expression levels were determined using the ΔΔCt method with the mRNA levels being normalized to the geometric mean of *PPIA* and *B2M*.

Primers or Taqman probes or Taqman expression assays used for mRNA levels measured in Fig. 2, Fig. 4 and Fig. 6 are shown in Supplementary Table 3.

#### DNA isolation and mtDNA quantification

Total DNA was extracted from ∼20 mg mouse gastrocnemius muscle using the Dneasy Blood and Tissue Kit (Qiagen) according to the manufacturer’s instructions and treated with RNase A. Levels of mtDNA were measured by qPCR using 2.5 ng of DNA in a QuantStudio 6 Flex Real-Time PCR System using TaqMan™ Universal Master Mix II, with UNG. The mt-Nd1 and mt-Nd6/Nd5 TaqMan gene expression assays were used. The 18S probe was used for normalization.

#### Aerobic and strength ET intervention in healthy young women

A detailed description of the study participants, research design and methods has been previously published^61^. In the present study, we included vastus lateralis muscle lysate for immunoblotting obtained from 9 healthy young women (age, 33 ± 6 years; body mass index (BMI), 23.2 ± 2.6 kg m^-2^) before and after 14-week of aerobic and strength exercise training.

#### High-intensity interval training intervention in healthy young men

A detailed description of the study participants, research design and methods has been previously published^63^. In the present study, we included vastus lateralis muscle lysate for immunoblotting obtained from 6 healthy young men before and after 6-week HIIT training.

#### Progressive resistance exercise training in male young and older individuals

A detailed description of the study participants, research design and methods has been previously published ^45^. The original study reported no additive effect of vitamin D intake during the 12 weeks of resistance exercise training on muscle hypertrophy or muscle strength ^60^. Accordingly, in the present study, samples from young or older participants were considered as one group, irrespective of vitamin D intake. We included vastus lateralis muscle samples (∼10 mg wet weight) for immunoblotting obtained from a subset of the original sample set due to lack of sample material. For the young participants, we included 10 samples before and 6 samples after resistance exercise training. For the older participants we included 11 before and 9 samples after resistance exercise training.

#### High-intensity interval training in middle-aged male individuals, who were lean, obese or obese with type 2 diabetes

A detailed description of the study participants, research design and methods has been previously published^62^. In short, 15 middle-aged men with T2D and obesity, 15 age-matched glucose-tolerant men with obesity, and 18 age-matched glucose-tolerant lean men were recruited. All participants, except four (two men with T2D and obesity and two lean men) completed the study. The training protocol consisted of 8-weeks with three weekly training sessions consisting of supervised HIIT in combination with biking and rowing. HIIT-sessions consisted of 5 x 1 min exercise blocks interspersed with 1 min rest, and shifted between blocks on cycle and rowing ergometers. The volume was increased from two to five blocks during the 8 weeks. In the present study, we included vastus lateralis muscle samples that were obtained 4-5 days after the last HIIT session but 48 hours after the last physical activity (a VO_2_ max test). The number of samples included for immunoblotting as follows: 16 before and after HIIT from lean glucose-tolerant men,15 before and after HIIT from glucose-tolerant men with obesity, and 13 before and after HIIT from men with T2D and obesity.

#### Ethical committee approval

Clinical experiments were approved by the Ethics Committee of Copenhagen or by the Regional Scientific Ethical Committees for Southern Denmark and performed in accordance with the Helsinki Declaration II. Studies including human study participants are described in Ref^60–63^. All mouse experiments complied with the European Convention for the protection of vertebrate animals used for experimental and other scientific purposes (No. 123, Strasbourg, France, 1985; EU Directive 2010/63/EU for animal experiments) and were approved by the Danish Animal Experimental Inspectorate (License number: 2016-15-0201-01043).

#### Graphical illustrations

Graphical illustrations were created in ©BioRender - biorender.com, as indicated.

#### Statistical methods

Statistical analyses were performed using GraphPad Prism 9 (version 9.3.1). Results are presented as mean ± SEM with individual values shown, when feasible. Statistical differences were analyzed by ordinary one-way ANOVA, repeated/ordinary two-way ANOVA, or Log-rank (Mantel-Cox) test, and Mann-Whitney test as applicable. Dunnett’s multiple comparisons test or Šídák’s multiple comparisons test were used to evaluate significant interactions in ANOVAs. As statistical tests varied according to the dataset being analyzed, the respective tests utilized are specified within the figure legends. 0.05≤p<0.1 were considered a tendency and p values <0.05 were considered significant.

## Acknowledgements

We acknowledge the technical assistance of Betina Bolmgren and Irene Nielsen, Martin Thomassen, and Roberto Meneses-Valdes (Department of Nutrition, Exercise and Sports, Faculty of Science, University of Copenhagen, Denmark), Anja Jokipii-Utzon (Institute of Sports Medicine, Bispebjerg Hospital, Copenhagen, Denmark), and Michala Carlsson (Department of Biomedical Sciences, University of Copenhagen, Denmark). We thank Vivian Shang for her assistance with the *Drosophila* experiments (Charles Perkins Centre, The University of Sydney, Australia). We acknowledge Cristiano di Benedetto for preparing muscle samples for TEM analysis (Core Facility for Integrated Microscopy, Faculty of Health and Medical Sciences, University of Copenhagen, Denmark).

## Funding

The study was supported by the Novo Nordisk Foundation (grant NNF16OC0023418, NNF18OC0032082, and NNF20OC0063577 to L.S.; grant NNF22OC0074110 to A.M.F.), by Independent Research Fund Denmark to L.S. (#0169-00013B), by the European Union’s Horizon 2020 research and innovation programme (Marie Skłodowska-Curie grant agreement No 801199 to T.C.P.P. and E.A.R.), and grants by Danish Council for Independent Research - Medical Sciences (4181-00078) and the Augustinus Foundation to H.P.

## CRediT authorship contribution statement

**Tang Cam Phung Pham**: Conceptualization, Methodology, Validation, Formal analysis, Investigation, Writing - Original Draft, Visualization, Project administration, Funding acquisition; **Steffen Henning Raun:** Conceptualization; Investigation, Writing - Review & Editing; **Essi Havula**: Methodology, Investigation, Writing - Review & Editing; **Carlos Henriquez-Olguín:** Methodology, Investigation, Writing - Review & Editing; **Diana Rubalcava-Gracia:** Methodology, Investigation, Writing - Review & Editing; **Emma Frank:** Methodology, Investigation, Writing - Review & Editing; **Andreas Mæchel Fritzen:** Methodology, Investigation, Writing - Review & Editing; **Paulo Jannig:** Methodology, Investigation, Writing - Review & Editing; **Nicoline Resen Andersen:** Investigation, Writing - Review & Editing; **Rikke Kruse:** Investigation, Writing - Review & Editing; **Mona Sadek Ali:** Investigation, Writing - Review & Editing; **Jens Frey Halling:** Resources, Writing - Review & Editing; **Stine Ringholm:** Resources, Writing - Review & Editing; **Solvejg Hansen:** Resources, Writing - Review & Editing; **Anders Krogh Lemminger:** Resources, Writing - Review & Editing; **Maria Houborg Petersen:** Resources, Writing - Review & Editing; **Martin Eisemann de Almedia:** Resources, Writing - Review & Editing; **Thomas E. Jensen:** Resources, Writing - Review & Editing; **Bente Kiens:** Resources, Writing - Review & Editing; **Morten Hostrup:** Resources, Writing - Review & Editing; **Steen Larsen:** Methodology, Investigation, Writing - Review & Editing; **Niels Ørtenblad:** Resources, Writing - Review & Editing; **Kurt Højlund:** Resources, Writing - Review & Editing; **Peter Schjerling:** Methodology, Investigation, Resources, Writing - Review & Editing; **Michael Kjær:** Resources, Writing - Review & Editing; **Jorge Ruas:** Resources, Writing - Review & Editing; **Aleksandra Trifunovic:** Resources, Writing - Review & Editing; **Jørgen Wojtaszewski;** Resources, Writing - Review & Editing; **Joachim Nielsen:** Methodology, Investigation, Writing - Review & Editing; **Klaus Qvortrup:** Resources, Writing - Review & Editing; **Henriette Pilegaard:** Resources, Writing - Review & Editing; **Erik Arne Richter:** Resources, Investigation, Writing - Review & Editing, Supervision, Funding acquisition; **Lykke Sylow**: Conceptualization, Methodology, Validation, Formal analysis, Investigation, Writing - Original Draft, Project administration, Supervision, Funding acquisition.

---

**Supplementary Fig. 1.**
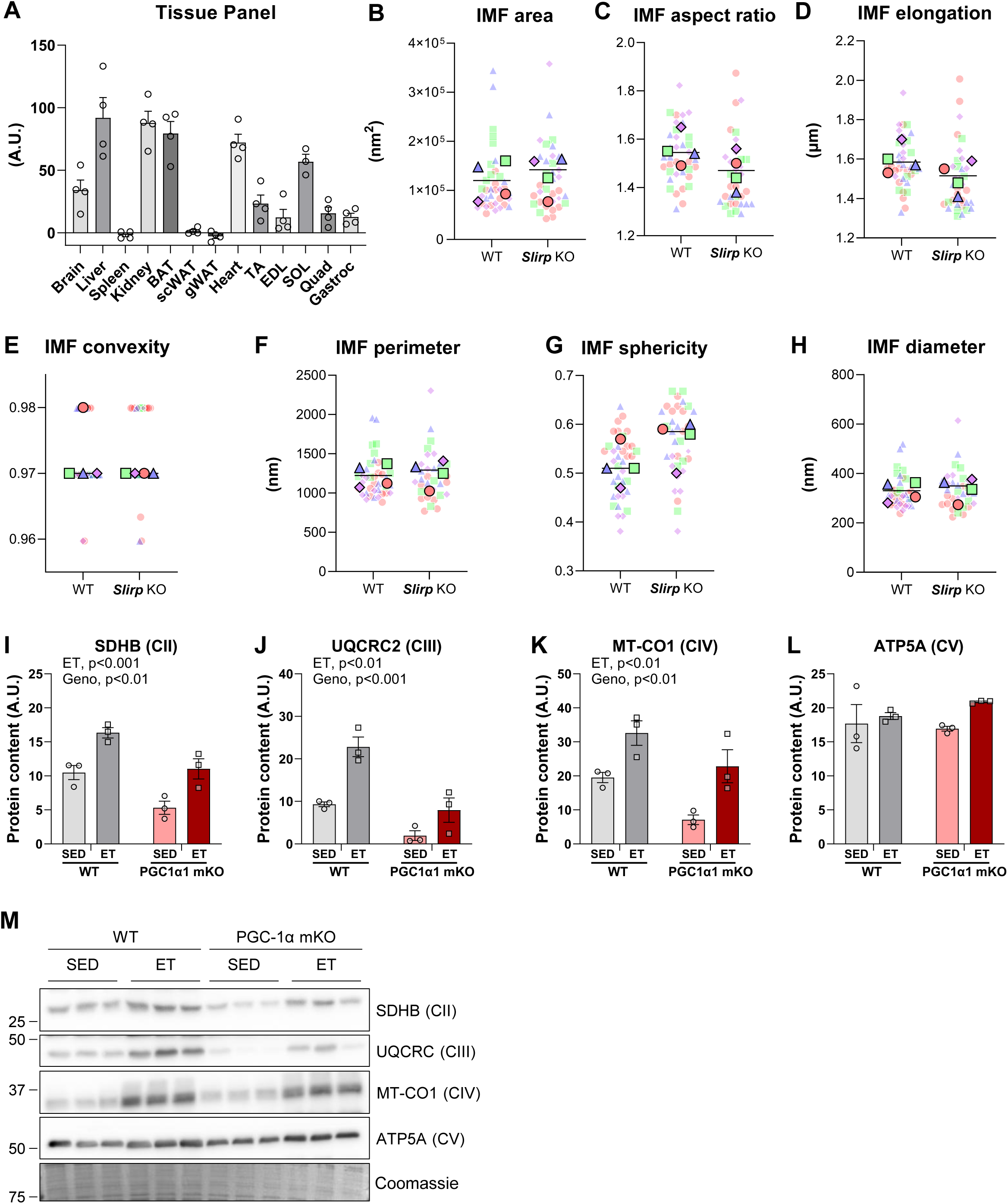
(A) Quantification of SLIRP protein content across different WT tissues (WT samples of n=4, 12 weeks of age) assessed by immunoblotting. (B-H) Transmission electron microscopy quantification of the following parameters of mitochondrial area in the intermyofibrillar (IMF) region (B), aspect ratio (C), elongation (D), convexity (E), perimeter (F), sphericity (G), and diameter (H). Small circles refer to each fiber, large squares in similar color refer to average of fibers per biological replicate, (female, n=4). (I-M) Western blot analysis of SDHB (CII), UQCRC2 (CIII), MT-CO1 (CIV), and ATP5A (CV) protein abundance in quadriceps of sedentary (SED) and 12-week exercise-training (ET) PGC-1α mKO and control littermates (WT) mice (subset of total cohort, male, n=3). Representative blots. Coomassie was used as loading control. Data are means ± SEM, including individual values. ET, main effect of exercise training; Geno, main effect of genotype. *p<0.05, **p<0.01, ***p<0.001, as per Unpaired Student’s t test (B-D, F-H), Mann Whitney test (E), Two-way ANOVA (I-L)

**Supplementary Fig. 2.**
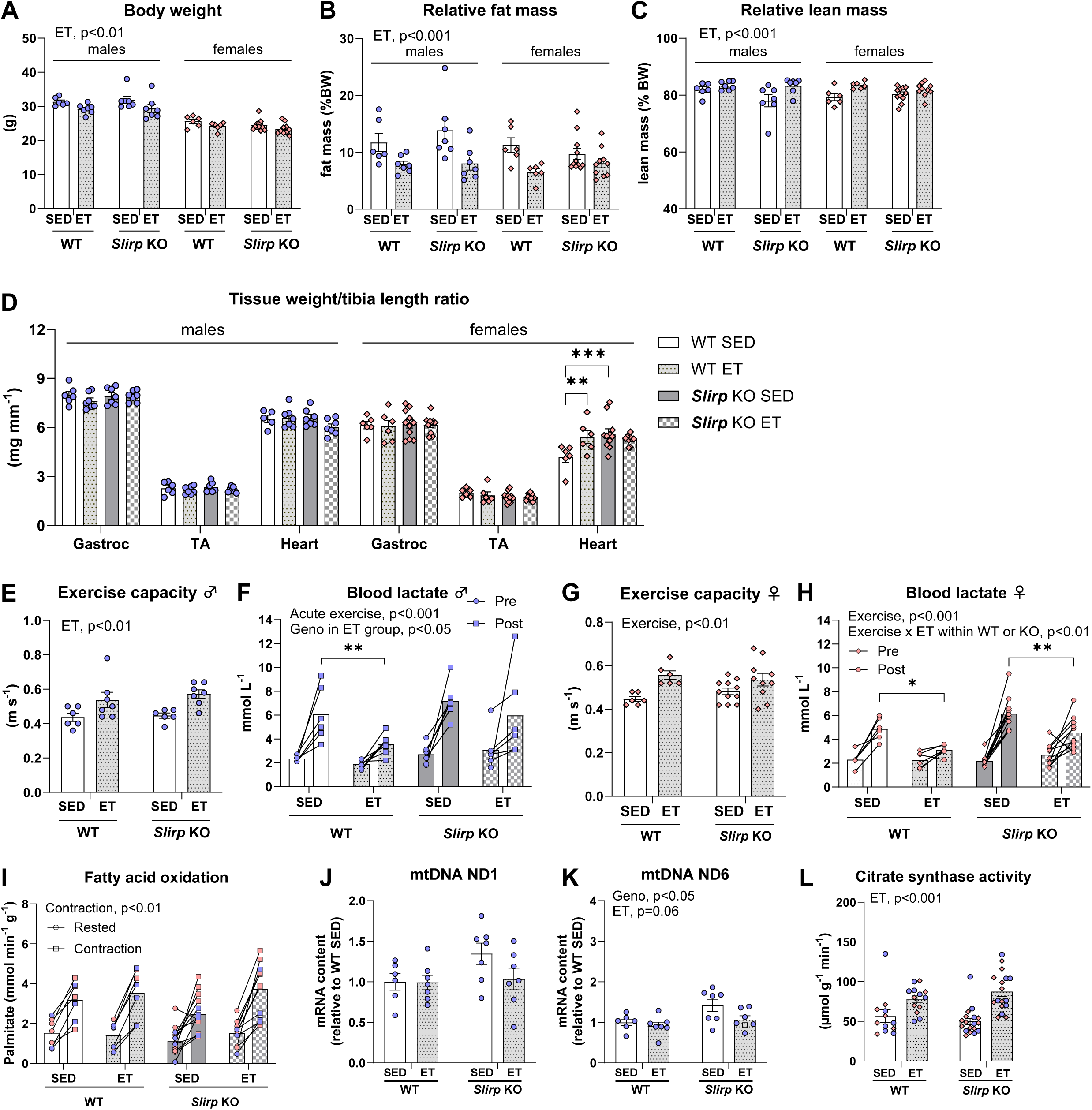
(A-C) Body weight, fat mass relative to body weight, lean mass relative to body weight of male and female SED and ET *Slirp* KO or WT mice after 6-weeks of ET (n=6-11/group, **○** (blue) male, ◇ (red) female).(D) Gastrocnemius, tibialis anterior (TA) and heart weight relative to tibia length of female SED and ET *Slirp* KO or WT mice after 10-weeks of ET (n=6-11/group, **○** (blue) male, ◇ (red) female).(E-H) Exercise capacity of male and female SED and ET *Slirp* KO or WT mice after 8-weeks of ET and corresponding blood lactate levels before and after exercise bout (n=6-11/group, **○** (blue) male, ◇ (red) female).(I) Fatty acid oxidation in isolated soleus of male and female SED and ET *Slirp* KO or WT mice after 10-weeks of ET. WT SED male/female, n=3/4, WT ET male/female, n=4/3, KO SED male/female, n=4/9, KO ET male/female, n=4/7. SED data is also depicted in Fig. 1H.(J-L) RT-qPCR analysis of mtDNA levels and citrate synthase activity in gastrocnemius of SED and ET *Slirp* KO and WT mice (n=6-11/group, **○** (blue) male, ◇ (red) female).Data are means ± SEM, including individual values where applicable. ET, main effect of exercise training; Geno, main effect of genotype; Acute exercise, main effect of acute exercise; Geno in ET group, main effect of genotype within exercise training group; Contraction, main effect of contraction.*p<0.05, **p<0.01, ***p<0.001, as per Two-way ANOVA within sex (A-D), Two-way ANOVA (E, G, J-L), Two-way RM ANOVA with Šídák’s multiple comparisons test (F, H, I).

**Supplementary Fig. 3.**
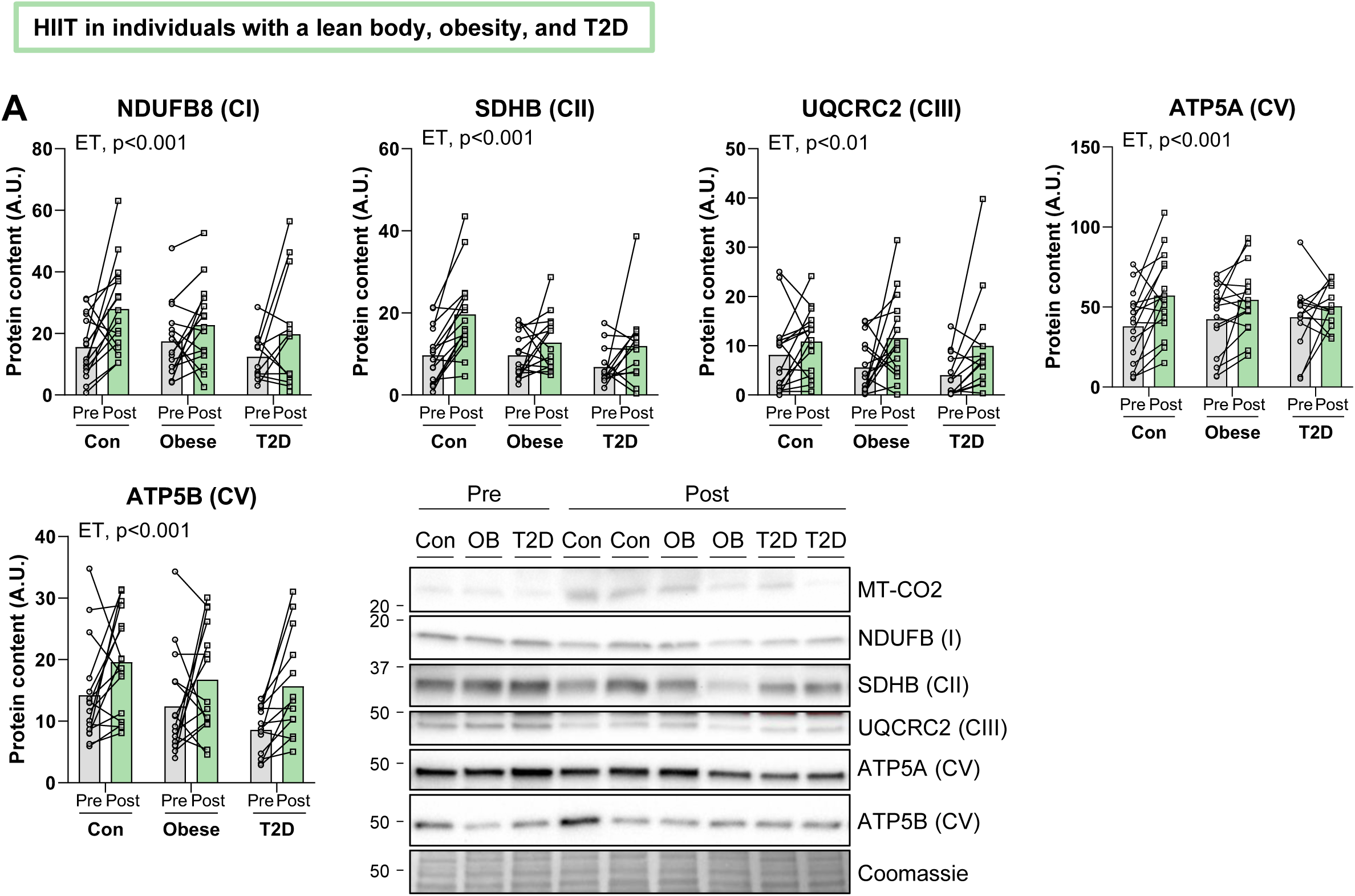
(B) Western blot analysis of NDUFB8 (CI), SDHB (CII), UQCRC2 (CIII), ATP5A (CV) and ATP5B (CV) protein in vastus lateralis of male individuals, who were lean (pre/post, n=16/16), obese (pre/post, n=15/15) or type 2 diabetic (pre/post n=13/13) after high-intensity interval training^62^, and representative images. Coomassie was used as control. Data presented as individual before-and-after values. ET, main effect of exercise training. *p<0.05, **p<0.01, ***p<0.001, Two-way RM ANOVA (A).

**Supplementary Table 2:**
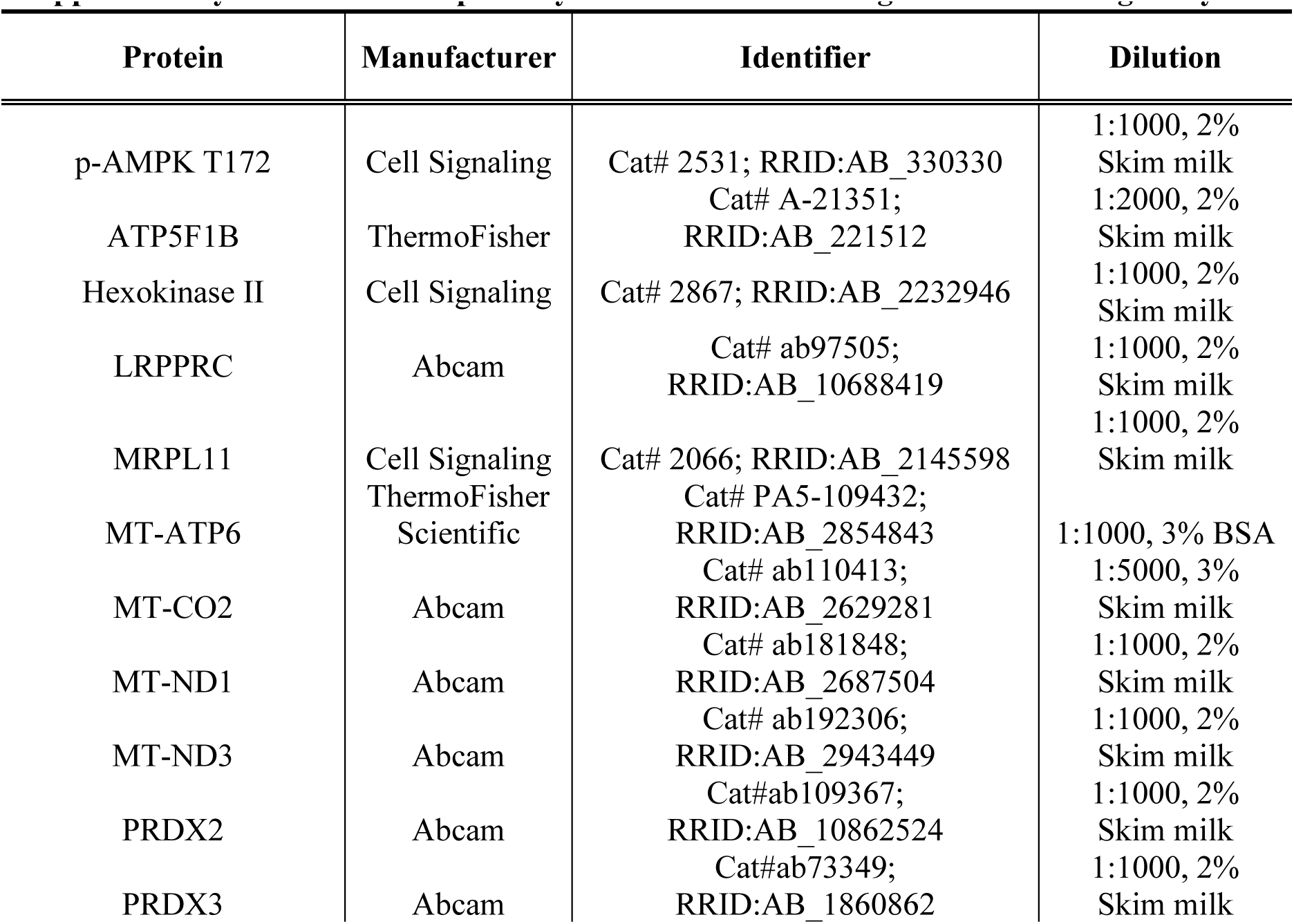
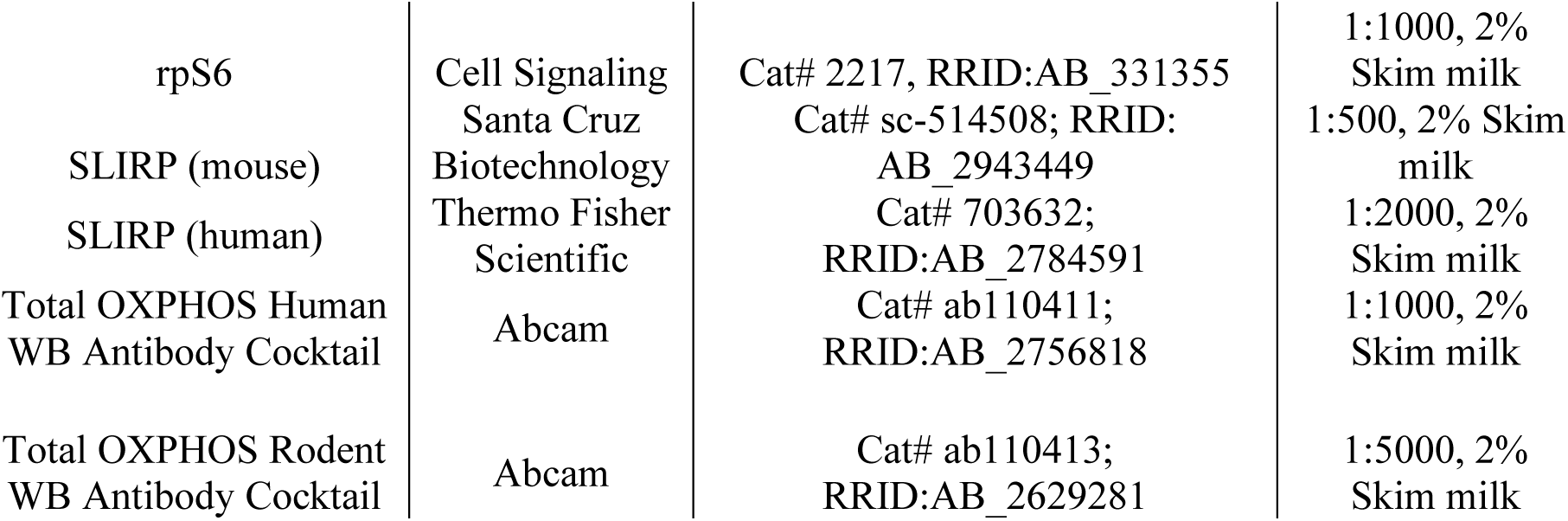
List of primary antibodies used during Western blotting analysis.

| Protein | Manufacturer | Identifier | Dilution |
| --- | --- | --- | --- |
| p-AMPK T172 | Cell Signaling | Cat# 2531; RRID:AB_330330 | 1:1000, 2% Skim milk |
| ATP5F1B | ThermoFisher | Cat# A-21351; RRID:AB_221512 | 1:2000, 2% Skim milk |
| Hexokinase II | Cell Signaling | Cat# 2867; RRID:AB_2232946 | 1:1000, 2% Skim milk |
| LRPPRC | Abcam | Cat# ab97505; RRID:AB_10688419 | 1:1000, 2% Skim milk |
| MRPL11 | Cell Signaling | Cat# 2066; RRID:AB_2145598 | 1:1000, 2% Skim milk |
| MT-ATP6 | ThermoFisher Scientific | Cat# PA5-109432; RRID:AB_2854843 | 1:1000, 3% BSA |
| MT-CO2 | Abcam | Cat# ab110413; RRID:AB_2629281 | 1:5000, 3% Skim milk |
| MT-ND1 | Abcam | Cat# ab181848; RRID:AB_2687504 | 1:1000, 2% Skim milk |
| MT-ND3 | Abcam | Cat# ab192306; RRID:AB_2943449 | 1:1000, 2% Skim milk |
| PRDX2 | Abcam | Cat#ab109367; RRID:AB_10862524 | 1:1000, 2% Skim milk |
| PRDX3 | Abcam | Cat#ab73349; RRID:AB_1860862 | 1:1000, 2% Skim milk |
| rpS6 | Cell Signaling | Cat# 2217, RRID:AB_331355 | 1:1000, 2% Skim milk |
| SLIRP (mouse) | Santa Cruz Biotechnology | Cat# sc-514508; RRID: AB_2943449 | 1:500, 2% Skim milk |
| SLIRP (human) | Thermo Fisher Scientific | Cat# 703632; RRID:AB_2784591 | 1:2000, 2% Skim milk |
| Total OXPHOS Human WB Antibody Cocktail | Abcam | Cat# ab110411; RRID:AB_2756818 | 1:1000, 2% Skim milk |
| Total OXPHOS Rodent WB Antibody Cocktail | Abcam | Cat# ab110413; RRID:AB_2629281 | 1:5000, 2% Skim milk |

**Supplementary Table 3:**
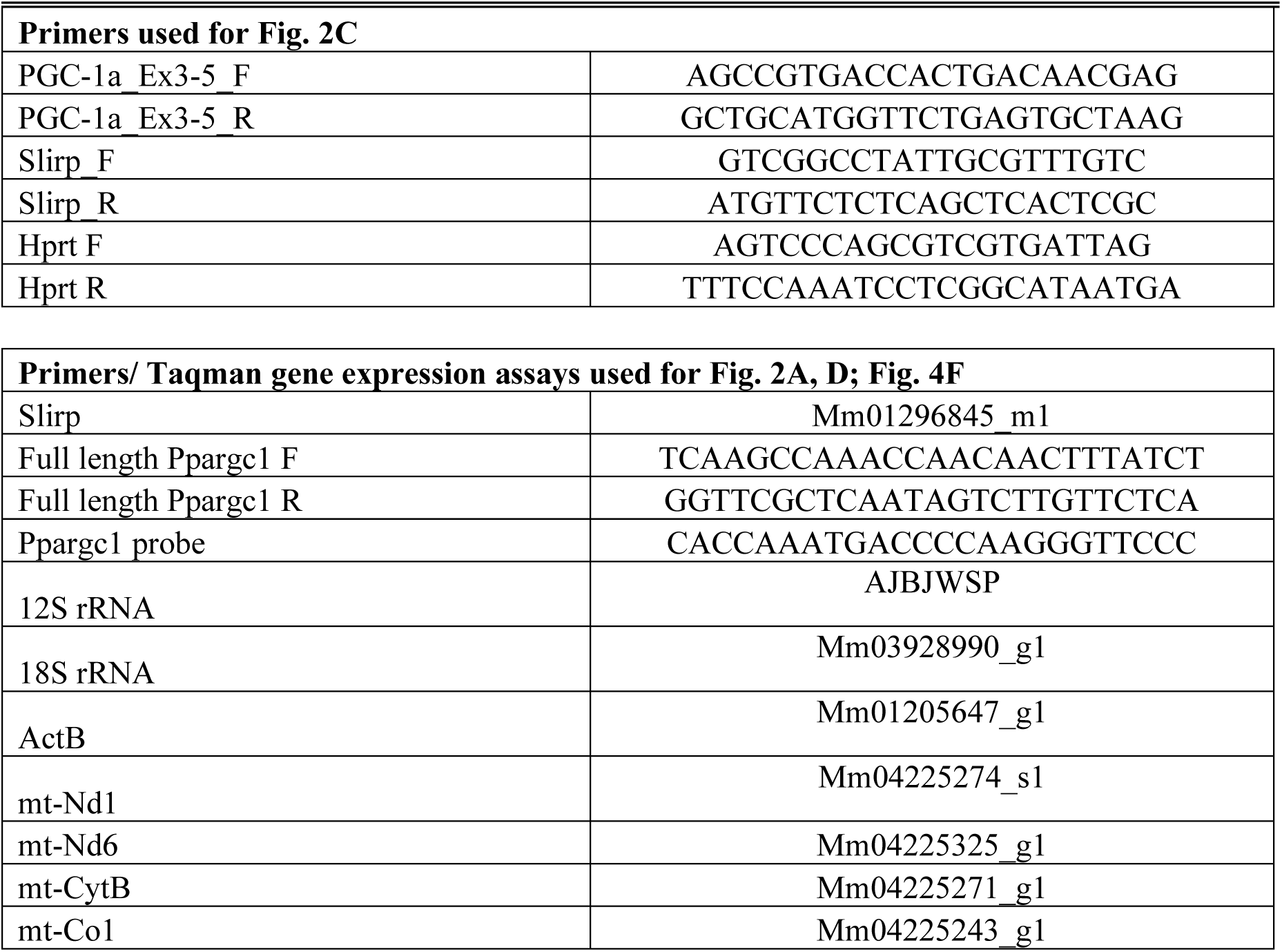
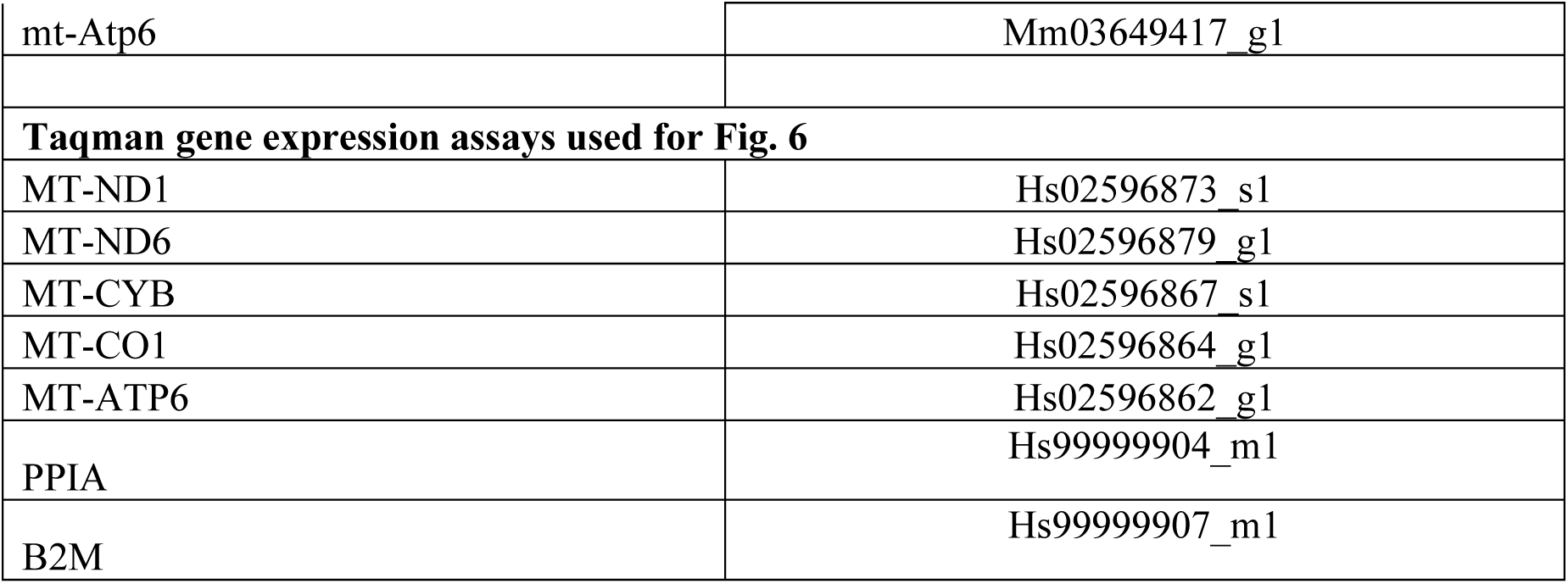
List of primers or Taqman probes used for RT-qPCR.

## Notes

### Competing Interest Statement

The authors have declared no competing interest.

### Summary of Updates

Fig 4. and Fig. 6 revised.

